# Genomic divergence and differential gene expression between crustacean ecotypes across a marine thermal gradient

**DOI:** 10.1101/2020.09.04.282517

**Authors:** Arsalan Emami-Khoyi, Ingrid S. Knapp, Daniela M. Monsanto, Bettine Jansen van Vuuren, Robert J. Toonen, Peter R. Teske

**Affiliations:** Centre for Ecological Genomics and Wildlife Conservation, Department of Zoology, University of Johannesburg, Auckland Park 2006, South Africa; Hawai‘i Institute of Marine Biology, University of Hawai‘i at Mānoa, Kāne‘ohe, Hawai‘i, Honolulu, HI, USA

**Keywords:** climate change, decapod crustaceans, ecological speciation, gene expression, genomic divergence, Pool-Seq, RNAseq, temperature adaptation, transcriptomics

## Abstract

Environmental gradients between marine biogeographical provinces separate distinct faunal communities; in the absence of absolute dispersal barriers numerous species nonetheless occur on either side of such boundaries. While the regional populations of such widespread species tend to be morphologically indistinguishable from each other, genetic evidence suggests that they represent unique ecotypes, and likely even cryptic species, that may be uniquely adapted to their local environment. Here, we explored genomic divergence in four sympatric southern African decapod crustaceans whose ranges span the boundary between the cool-temperate west coast (south-eastern Atlantic) and the warm-temperate south coast (south-western Indian Ocean) near the southern tip of the African continent. Using genome-wide data, we found that all four species comprise distinct west- and south coast ecotypes, with molecular dating suggesting divergence during the Pleistocene. Using transcriptomic data from one of the decapod crustaceans, we further found a clear difference in gene expression profiles between the west- and south coast ecotypes. This was particularly clear in the individual from the south coast, which experienced a ‘transcriptomic shock’ at low temperatures that are more typical of the west coast and may explain their absence from that coastline. Our results shed new light on the processes involved in driving genomic divergence and incipient speciation.

## Introduction

The assumption that populations of marine species are ‘open’ and thus exhibit little genetic differentiation along vast geographical scales has been challenged by numerous studies. It is now generaly accepted that range distributions can be significantly constrained by oceanographic boundaries such as upwelling systems, currents, eddies and temperature gradients (Wilson et al. 2013). However, the fact that these boundaries are in most cases unsuitable to completely isolate coastal habitats, but nonetheless define distinct species assemblages and genetically unique populations of more widespread species (Pelc et al. 2009; Teske et al. 2011; Haye et al. 2014), has generated an interest in identifying environmental drivers of adaptation that may reduce the fitness of migrants once they disperse across these boundaries, and thus physiologically or competitively exclude them from adjacent marine regions (Sanford et al. 2003; Schmidt et al. 2008). Such information may contribute towards gaining a better understanding of the role of ecological factors in driving biodiversity generation.

Here, we explore temperature-driven population divergence across a thermal gradient along the south-western African coastline. This region comprises two major marine biogeographical provinces, the cool-temperate Namaqua Province on the west coast and the warm-temperate Agulhas Province on the south coast, and an intermediate area of faunal overlap on the south-west coast that is sometimes treated as a distinct province (Lombard et al. 2004). The cool-temperate and warm-temperate provinces are not isolated from each other: particularly during El Niño events, warm-water species from the east may temporarily settle on the west coast in large numbers (Branch 1984; Roy et al. 2001; Dufois and Rouault 2012). Despite these moderate levels of connectivity, widespread species are often subdivided into genetically and sometimes morphologically distinguishable sister populations whose ranges rarely extend far beyond the boundaries of a specific biogeographical province (Teske et al. 2011)

Genetic structure of coastal animals in southern Africa has primarily been studied using mitochondrial DNA (mtDNA) sequence data, but recent studies have highlighted numerous limitations of this marker in understanding the evolution of regional populations. Specifically, mtDNA data do not support congruent patterns in all of the species studied, with some displaying deep phylogenetic splits and others being genetically homogeneous (Teske et al. 2011). This discrepancy has been attributed to limitations in the power of mtDNA to detect recent divergence, as phylogenetic splits in supposedly non-structured species were subsequently identified by means of nuclear introns or larger genomic datasets (Teske et al. 2014; Teske et al. 2019).

In the present study, we used genomic data to explore whether the discrepancies between species divergence using mtDNA could be an artefact of the insufficient resolution, as previously suggested by Teske et al. (2019). We further aimed to determine whether west coast and south coast populations of co-distributed decapod species diverged from their respective common ancestors at approximately the same time, which would indicate that the same historical event was responsible for driving incipient speciation. In addition, we used transcriptomic data of one of the study species to assess whether individuals from the west and south coast respond differently to cold temperature stress and investigate whether adaptive differences drive early-stage ecological speciation.

## Methods

### Genomic analyses

Genomic data from pooled population samples were generated to determine whether divergence between west coast and south coast populations of co-distributed decapod crustaceans occurred at approximately the same time. To this end, data were generated from four species that are abundant in estuarine habitats located on the west and south coast of South Africa. These were the sand prawn *Callichirus kraussi* (Stebbing, 1900), the mudprawn *Upogebia africana* (Ortman, 1894), the crown crab *Hymenosoma orbiculare* Desmarest, 1823 and the hermit crab *Diogenes brevirostris* Stimpson, 1858 (Fig. 1, APPENDIX A – Table A1).

**Figure 1.**
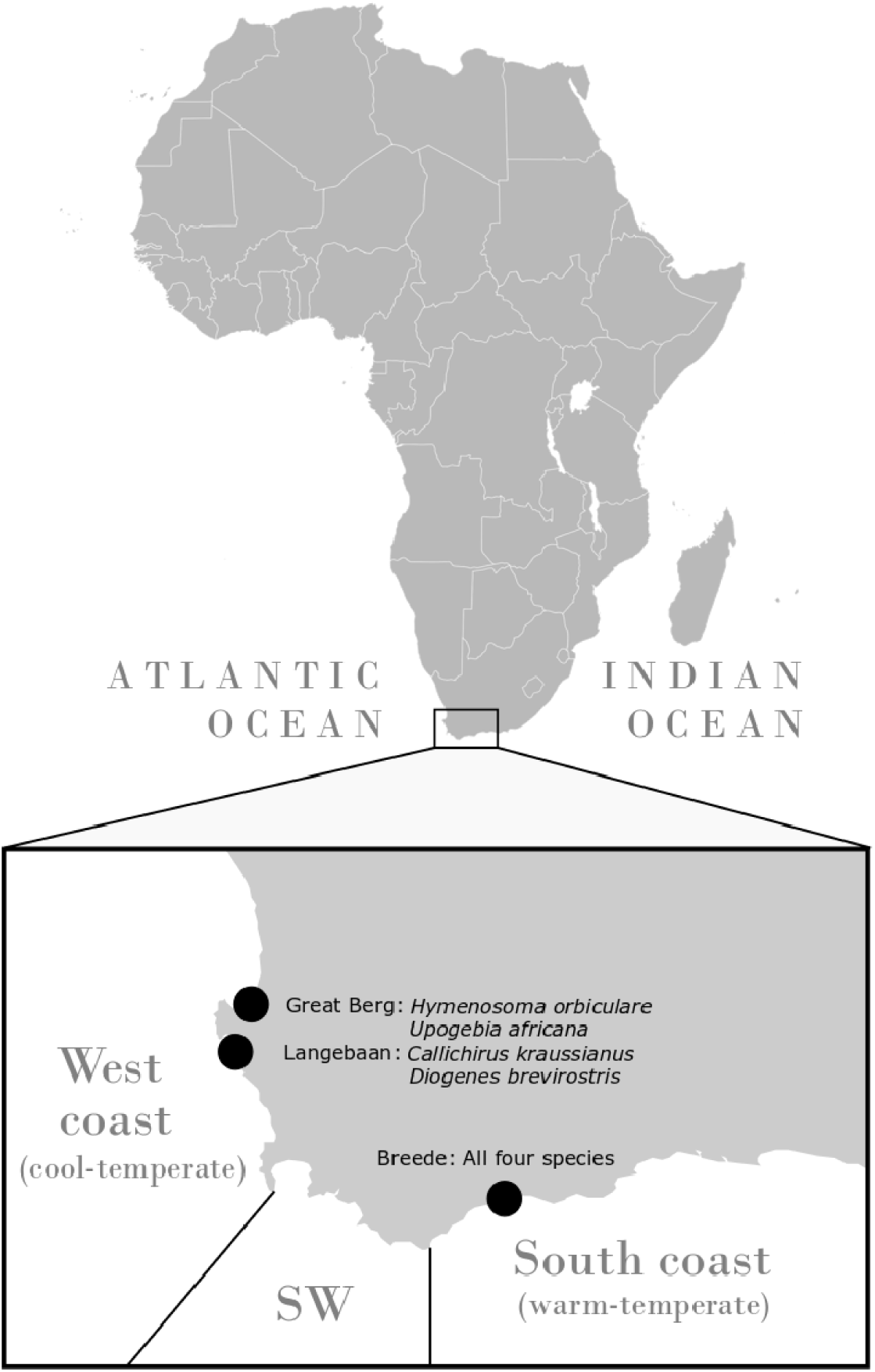
Map of the sampling area at the southern tip of Africa, showing the location of sampling sites and the species collected at each site. West coast sites are in the Atlantic Ocean and south coast sites are in the Indian Ocean. The southwest coast (SW) was not sampled because of previous evidence for introgression in this environmentally intermediate area.

#### Generation of raw sequence data

Genomic DNA of high molecular weight was extracted from muscle tissue of the study species using the CTAB protocol (Doyle and Doyle 1987) and the quality of the extractions was assessed using a NanoDrop™ 2000c spectrophotometer (Thermo-Fisher Scientific, Waltham, MA, USA). The genomic libraries were prepared from the pool of DNA extractions following the ezRAD protocol (Toonen et al. 2013) using DPNII, a frequent-cutter restriction enzyme (New England Biolabs, Ipswich, MA, USA) in a 50 μl reaction (1 μl of 1x DPNII, 5 μl of 10x buffer 3.1, and 44 μl of DNA extraction and HPLC grade water) and incubated at 37 °C for 3 hours, 20 minutes at 65 °C and then held at 15 °C. Following enzymatic digestion, samples were cleaned using Agencourt AMPure XP beads (Beckman Coulter, Indianapolis, IN, USA) at a DNA: beads ratio of 1:1.8. Library preparation was completed using the KAPA HyperPrep kit (Roche, Indianapolis, IN, USA) following manufacturer’s protocols including a size selection step with SPRIselect beads (Beckman Coulter, Indianapolis, IN, USA) to only retain 300–700 base pair (bp) fragments, and a library amplification step following the recommended cycle numbers to obtain approximately 1 μg of Illumina TruSeq adapter-ligated genomic library. The completed libraries were then passed through quality control steps (bioanalyzer and qPCR) prior to high-throughput sequencing by the Hawai□i Institute of Marine Biology Genetics Core Facility on the Miseq genomic analyzer platform (Illumina, San Diego, CA, USA) using the Illumina v3.2 kit with 300 bp paired-end chemistry.

#### Quality filtering of raw data, alignment and variant calling

The quality of raw sequences was checked using the FastQC v0.11.1.5 package (Andrews 2017). Leading and trailing low-quality bases (with Phred score < 3), sliding 4-bp windows with average Phred score < 20 Phred, and Illumina adapter contaminations were removed using Trimmomatic v0.36 (Bolger et al. 2014). In the absence of reference genomes for the four non-model decapod crustaceans, the annotated genome of the Chinese mitten crab, *Eriocheir sinensis* (Song et al. 2016), was used as a surrogate reference genome against which quality-filtered sequences were mapped using the semi-global aligner BWA-MEM v0.7.12 (Li 2013). As mitochondrial DNA often violates the expectations of the neutral theory of molecular evolution (Kimura 1979) required for accurate molecular dating and tends to be subject to strong selection (Ballard and Kreitman 1995; Meiklejohn et al. 2007; Stewart et al. 2008), mitogenome sequences were excluded from the mitten crab’s draft assembly by conducting a BLAST search of the mitten crab mitogenome (NCBI accession number NC_011598.1) against the assembled mitten crab scaffolds, and removing scaffolds that originated from mitochondrial DNA sequences. Conversion and sorting of sequence alignment map (SAM) files to BAM format were done with Samtools v1.6 (Li et al. 2009). Only sequences with a minimum alignment quality score of ≥ 40 Phred (probability of erroneous alignment ≤ 10^−4^) were selected. This resulted in unmapped sequences and sequences that were mapped to multiple positions in the mitten crab draft genome to be excluded from downstream variant calling.

A combination of Samtools v1.6 mpileup and VarScan v2.3.9 mpileup2snp commands (Koboldt et al. 2013) was used to find variant sites for each species. In VarScan, the minimum read coverage to call a variant site was set to > 4 reads, the p-value was set to 0.01, and the minimum variant allele frequency was set to a number equal to 0.5 divided by the number of individuals in each pool (Koboldt et al. 2013). All indel sites and variants with > 90% support on one strand (forward or reverse reads) were removed.

Variant sites that are polymorphic in individuals should be polymorphic in the high coverage pool sequencing of a population. However, the absence of one allele in a population can arise due to uneven sequencing depth rather than fixation or loss that follows genetic drift, so only biallelic variant sites (i.e., single nucleotide polymorphisms or SNPs) that were genotyped in both southern and western populations were included in subsequent analyses.

#### Estimating effective population size from Pool-Seq data

Effective population size can be calculated using Ewens’ formula for estimating the scaled mutation rate under the infinite allele model (Ewens 1972), □ = 4*N*_*e*_µ, where *N*_*e*_ is the effective population size of organisms and µ is a neutral mutation rate per locus per generation. The estimation of *N*_*e*_ is possible by calculating an estimator of □ with a known mutation rate. Watterson’s estimation of □,*ŵ*(Watterson 1975) was selected here, which is based on the number of segregating sites in each population. It has been shown to be particularly suitable when the number of sequences is large and the majority of rare variants are genotyped. (RoyChoudhury and Wakeley 2010). A major challenge in Pool-Seq pipelines is to correctly estimate allele frequencies in pools of populations with a different number of individuals and sequencing depth (Kofler et al. 2011), and *ŵ* is potentially less sensitive to this methodological limitation than the more commonly used but more complex alternative, the Tajima’s 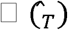 (Tajima 1983), which requires estimation of both the number of segregating sites and the estimation of allele frequencies from each pool of individuals.

Watterson’s estimator of □ was calculated in Popoolation v1.2.2 (Kofler et al. 2011) by applying a 1000 bp sliding window. The same program was used to calculate Tajima’s *D* (Tajima 1989) as a means of confirming that the majority of the SNP data conformed to the expectations under the neutral theory of molecular evolution (Kimura 1979). Estimation of population genomic statistics from pools of DNA is sensitive to uneven sequence coverage and the size of sliding windows (Kofler et al. 2011). In particular, genomic locations that demonstrate exceptionally high coverage are more prone to sequencing error. To address these issues, pileup files for each population were sub-sampled, without replacement, to an even sequence depth that did not exceed 50% of the sequence coverage range (maximum coverage - minimum coverage) across each population. Furthermore, sequence coverage for each sliding window was estimated, and approximately 30% of the sliding windows that were covered with the highest number of quality-filtered sequences were sub-selected to estimate Watterson □ and Tajima’s *D*. Popoolation2 v1201 (Kofler et al. 2011) was used to estimate allele frequency changes and *F* _ST_ for each SNP between southern and western populations, and Fisher’s exact test (Fisher 1922) was performed to test the significance in allele frequency changes between population pairs.

#### Divergence time estimation

KimTree2 (Clemente et al. 2018a) was used to estimate divergence times between the southern and western populations of each species on a diffusion time scale (*τ*_*i*_ = time in generation / 2*N*_*e*_). Briefly, KimTree2 applies a hierarchical Bayesian approach (Gautier and Vitalis 2012) to model allele frequency changes from the most ancestral population (root) forward to the terminal nodes based on a population tree that is defined *a priori*. The program implements Kimura’s time-dependent diffusion approximation for genetic drift model (Kimura 1964), which assumes that all observed polymorphisms in the current populations were present in the common ancestor. Kimura’s diffusion model predicts that genetic drift occurs independently in each branch of the tree, and in the absence of migration and recurrent mutation after divergence, the random changes in allele frequency from one generation to the next are thus proportionate to the length of branches leading to each population on a diffusion time-scale (*τ*_*i*_) (Kimura 1964). KimTree2 implements a Metropolis-Hastings with Gibbs sampler (Geman and Geman 1987) to sample the posterior distribution of the parameters of interest (in this case, the branch lengths) throughout the population tree. This makes it possible to analyze large genomic datasets that exceed the capabilities of alternative coalescent-based methods. The reason for this is that coalescence-based methods follow a probabilistic framework to calculate the likelihood of all unknown genealogies that are compatible with the observed sample of genes (Kingman 1982). Integrating such genealogies using Markov Chain Monte Carlo methods becomes computationally more extensive with larger sample size and the complexity of demographic dynamics (Marjoram and Tavaré 2006; Beaumont 2008). Moreover, unlike many coalescent-based methods, diffusion model approximations applied in KimTree2 can be used to analyze Pool-Seq data, thus benefitting from the latter approach’s high accuracy-to-cost ratio (Schlötterer et al. 2014). KimTree2 was run with an unascertained flag in pool sequencing mode for three independent chains of 3×10^11^ iterations following 1×10^11^ burn-in steps. The allele frequencies in the root were sampled from a fixed beta distribution with shape parameters (□, β) set to 0.7. The length and number of initial pilot chains were increased to 50 chains with 10,000 iterations each to reach a proposal acceptance rate of approximately 30% to 40%. Effective sample size (ESS), autocorrelation, and convergence were monitored using a combination of KimTree2 log files and the R package Coda v0.19-3 (Plummer et al. 2006). In Coda, convergence between independent chains for each set of simulations was tested using Gelman and Rubin’s shrinking factor (Gelman and Rubin 1992).

To convert the divergence time on a diffusion time scale to years (assuming a generation time of 1 year), the branch length for each population was multiplied by 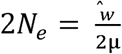, where μ is the median neutral mutation rate of 4.5 × 10^−9^ per site per generation of *Drosophila melanogaster* (Huang et al. 2016).

### Differential gene expression analyses

To assess whether exposure of the crustaceans to colder temperatures would produce different gene expression profiles between the west and south coast populations, we selected one of the four species, the mudprawn *Upogebia africana*, for differential gene expression analyses. We hypothesized that the species’ south coast population, which is much less likely to be subject to cold water upwelling than the west coast population, would produce a more significant transcriptomic response when exposed to colder temperatures due to thermal stress.

Six specimens each were obtained from the Great Berg Estuary (cool-temperature west coast) and the Breede Estuary (warm-temperate south coast), respectively. At the time of collection, temperatures in both estuaries were close to 20 °C. Specimens from each region were divided into two groups of three individuals each, which were kept in different, well-aerated aquaria at 20°C for an initial 24 hours to reduce the effect of acute stress responses following temperature change (Costas et al. 2011). Over the next 24 hours, the temperature in two aquaria (three individuals from the west coast and three from the south coast) was gradually decreased to 10 °C in increments of 1 °C every 2-3 hours. After completion of the experiment, prawns were killed immediately by means of a longitudinal scalpel cut through the centre-line of head, thorax and tail, and a portion of the hepatopancreas was excised and stored in RNA*later*™ solution (Thermo-Fisher Scientific, Waltham, MA, USA). Total RNA was extracted from each specimen using a combination of mechanical homogenization with Trizol and Qiagen RNeasy purification kit (Qiagen, Hilden, Germany). Then, cDNA libraries were constructed, indexed separately (Emami-Khoyi et al. 2020), and sequenced on an Illumina HiSeq™ 4000 platform (Illumina Inc., San Diego, USA) following the manufacturer’s instructions for 150 bp paired-end chemistry.

The reference hepatopancreas transcriptome of *U. africana* was assembled de-novo using the Trinity v2.8.6 pipeline (Haas et al. 2013). The longest reading frames within the assembled transcriptome were predicted in TransDecoder v5.5.0 (Haas and Papanicolaou 2016) and the final assembly was functionally annotated following the steps described in the Trinotate pipeline. In Trinotate, assembled transcripts were searched against multiple proteins and metabolic pathway entries deposited in the Swiss-Prot (https://www.uniprot.org/), NCBI non-redundant protein (https://www.ncbi.nlm.nih.gov/refseq/), and Kyoto Encyclopedia of Genes and Genomes (KEGG) (https://www.genome.jp/kegg/) databases. Protein domains for all transcripts were separately predicted in HMMER v3.1 (Finn et al. 2011) based on the homology to the known proteins in the Pfam (https://pfam.xfam.org/) database. In addition, transmembrane proteins and ribosomal RNA were identified using TmHMM v2 (Krogh et al. 2001) and RNAMMER v1.2 (Lagesen et al. 2007), respectively.

The expression levels of different genes were quantified using RSEM v1.3.1 (Li and Dewey 2011), and statistical significance in differences in expression profiles between different conditions (False Discovery Rate [FDR] < 0.05) were tested using empirical Bayes hierarchical model applied in the R package EBSeq v1.26.0 (Leng and Kendziorski 2014). The number of iterations in EBSeq was increased to 10 to achieved consistent results across multiple independent runs.

To identify major transcripts (genes and potential isoforms) that were differentially expressed between treatments, a subset of 225 transcripts with the highest levels of posterior fold changes, which corresponds to 2% and 4% of all genes that were differentially expressed at 10 °C compared to 20 °C, were selected, and their enrichment for different biological processes in terms of Gene Ontology categorizations (GO) (Botstein et al. 2000) was investigated using the g:Profiler online server and gProfiler2 R package. (Reimand et al. 2007) and applying g:SCS method for computing multiple testing correction for p-values. The ‘biological process’ category was selected since integrated fitness of biochemical pathways involved in the two other major categories (cellular component and molecular functions) will ultimately express itself in the form of biological processes that promote or prevent local adaptations. Moreover, enrichment analysis is sensitive to the selection of background gene sets against which the statistical significance for enrichment or depletion is tested. In crustaceans, several gene sets for different species of *Daphnia* are available. However, the genome size of these species are between 20% to 30% the size of the draft genome reported for the mitten crabs that was used as a surrogate genome in this study (Colbourne et al. 2011). The limited biochemical repertoire of the genome in *Daphnia* thus makes this genus, potentially unsuitable for the current study. As an alternative, the complete gene set from *Drosophila melanogaster* (version e97_eg44_p13_d22abce, 16/10/2019) was selected as the background gene set for enrichment analysis. In addition, the metabolic pathway content of the transcripts were predicted using the KEGG mapper online server (Kanehisa and Sato 2020), and the results were summarized in GAEV (Huynh and Xu 2018).

## Results

### Genomic analyses

The pool sequencing of the four decapod species produced 4,322,326 (*Callichirus kraussi*), 2,684,732 (*Upogebia africana*), 2,437,128 (*Diogenes brevirostris*) and 2,066,939 (*Hymenosoma orbiculare*) paired-end reads. The mean percentage of properly mapped reads varied from 91.35% in the southern population of *H. orbiculare* to 99.66% in the western population of *U. africana* (APPENDIX A – Table A1). The majority of identified variant sites were present in one population but not the other, and the total number of SNPs used in subsequent analyses was thus several orders of magnitude lower, ranging from 1,254 in *U. africana* to 2,829 in *H. orbiculare* (APPENDIX A – Table A2).

The mean *ŵ* throughout the genome was 0.0177 (95% CI: 0.007-0.026) for *C. kraussi*, 0.0234 (95% CI: 0.0120-0.0346) for *U. africana*, 0.004 (95% CI: 0.001-0.006) for *H. orbiculare*, and 0.020 (95% CI: 0.014-0.026) for *D. brevirostris*. The mean and mode of Tajima’s *D* estimated from each population were zero for all four species. This confirms that the vast majority of loci used in the analysis conformed to the neutral model of evolution. Furthermore, it confirmed that the selection of Watterson’s estimator of □ is unlikely to bias our results since in the neutrally evolving portions of the genomes, both estimations of □ ultimately converge on a unique value (Tajima 1989).

Fisher exact tests showed that allele frequencies between southern and western populations were significantly different (□ ≤ 0.05) in 3.23% of loci in *C. kraussi*, 2.42% of loci in *U. africana*, 4.17% loci in *H. orbiculare* and 2.05% of loci in *D. brevirostris*. The median of *F* _ST_ between southern and western populations ranged from 0.034 in *U. africana* to 0.0615 in *C. kraussi* (APPENDIX A – Table A3).

#### Estimation of branch lengths (*τ*_*i*_)

Gelman and Rubin’s shrinking factors and its 95% CI for *H. orbiculare* and *C. kraussi* were below the 1.1 convergence threshold (Brooks & Gelman, 1998). However, the upper 95% CI of shrinking factors in *U. africana* and *D. brevirostris* were slightly higher (< 1.4). Shrinking factors decreased to less than 1.1 after increasing the number of iterations to 7.5×10^11^, showing that convergence had been reached for all independent runs (Brooks and Gelman 1998) (APPENDIX A – Table A4). Examination of KimTree2 trace files confirmed that the burn-in and the total number of iterations were sufficient for all replicate runs to converge on similar values. Estimates of branch lengths (*τ*_*i*_) on the diffusion time-scale were slightly higher for the southern populations than the western populations and thus indicate slightly different levels of genetic drift between regions, an exception being *U. africana*, where the branches leading to the southern and western populations were approximately the same length (APPENDIX A – Table A5). After converting branch lengths (*τ*_*i*_) to time in years, the common ancestors of all species were dated to the Middle to Late Pleistocene. In each species, the 95% CIs for divergence time between pair of populations overlapped broadly (Fig. 2)

**Figure 2.**
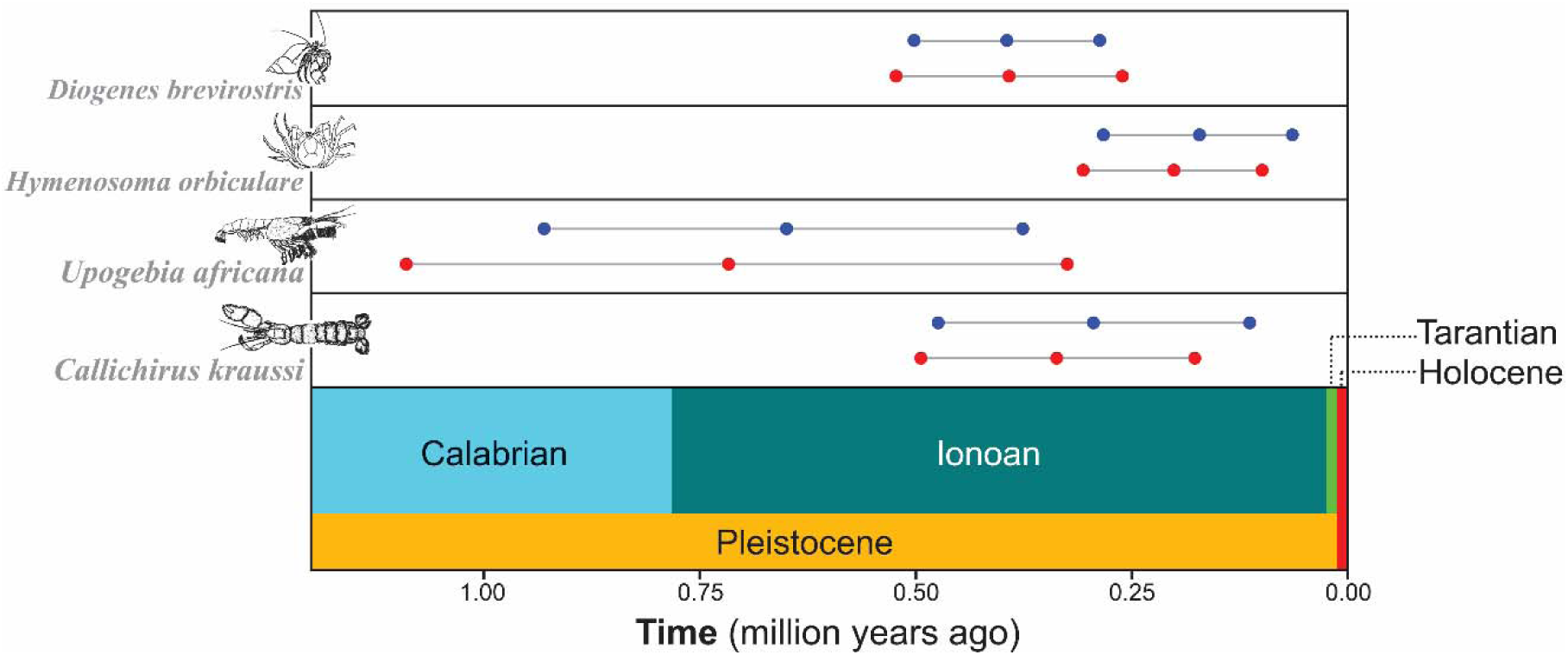
Time of divergence of the south coast (red) and west coast (blue) populations of four southern African decapod crustaceans from their respective ancestral populations estimated using KimTree2. Divergence time estimates (central circle) are flanked by circles that represent 95% confidence intervals. The y-axis depicts time in on Pleistocene and Holocene epoch.

### Differential gene expression analyses

#### Identification of differentially expressed (DE) genes in *Upogebia africana*

Trinity de-novo assembly of *Upogebia africana* produced 664,707 transcripts with a N50 of 1152 bp (APPENDIX A-TABLE A6). At 20 °C, the number of differentially expressed transcripts (< 0.05 FDR) between the west coast and the south coast site was 5,384, and this number increased to 9,639 in the specimens exposed to 10 °C (Fig. 3a), suggesting a “transcriptomic shock” in the south coast lineage at the lower temperature.

**Figure 3.**
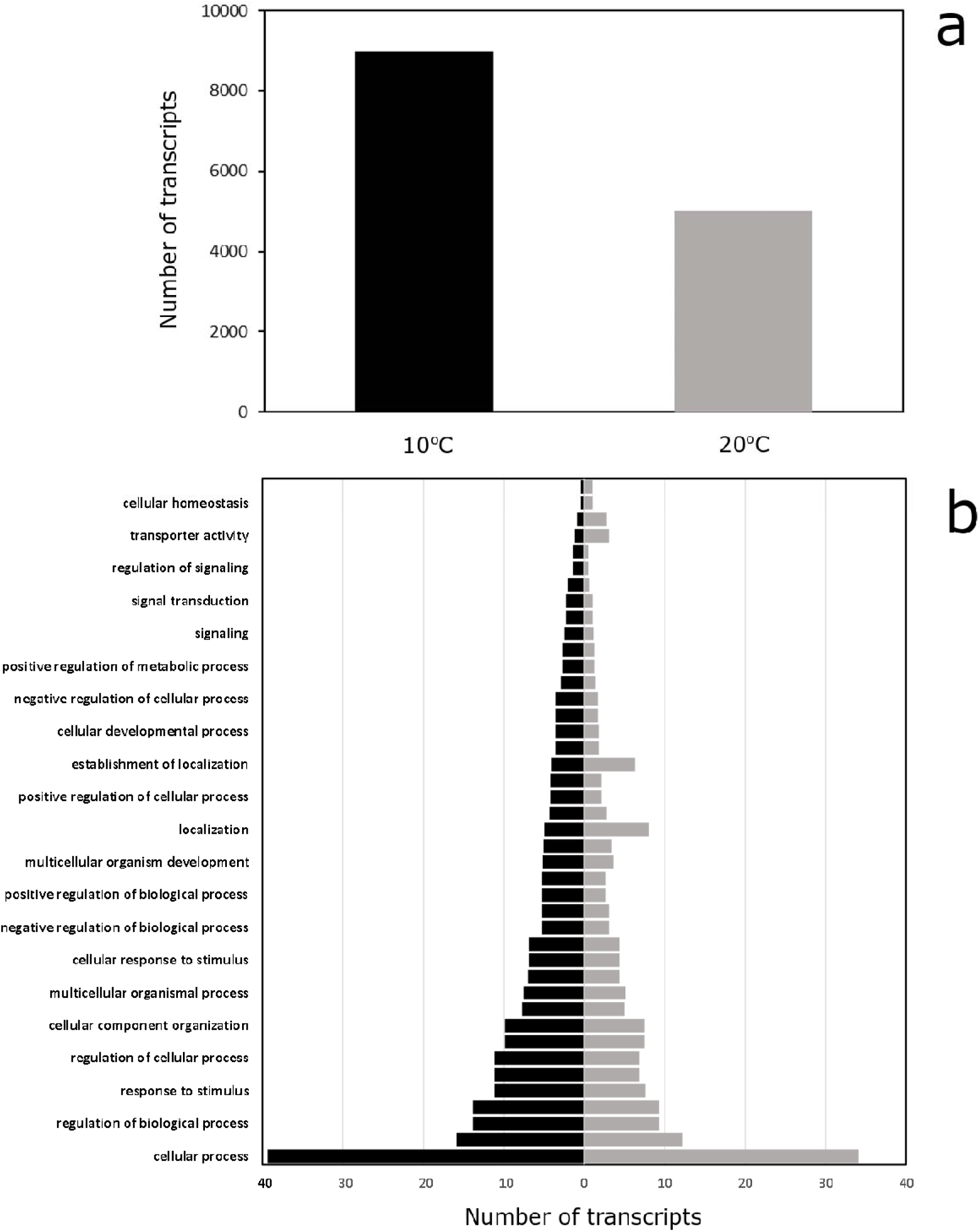
Total number of differentially expressed transcripts and the functional significance of 225 transcripts with highest posterior fold changes in term of Gene Ontology categorization biological process between west coast and south coast populations of *Upogebia africana*; a) comparison of the number of transcripts (genes and potential isoforms) that were differentially expressed (i.e., up- or downregulated) at a false discovery rate (FDR) < 0.05 in the south coast population compared to the west coast population at 10 °C (black) and 20 °C (grey), indicating a stronger transcriptomic response in the south coast population at colder temperatures (which are much more likely to be experienced by the west coast population); b) functional annotation of transcripts that were differentially expressed the two temperatures in terms of Gene Ontology biological process categorization.The raw data for generating this table is available at APPENDIX A-TABLES A7.

The majority of transcripts with the highest fold changes in abundance between treatments were involved in biological processes such as cellular processes, regulation of biological processes, and responses to external stimuli (Fig. 3b). Compared to the gene expression profile of control specimens kept at 20 °C, the expression profiles of individuals from the warm-temperate south coast that were exposed to 10 °C show a depletion in three major biological processes involved in responses to temperature stimuli (GO:0009266), immune effector processes (GO:0002252) and inflammatory responses to wounds or wound healing (GO:0042060, GO:0009611). These biological processes are involved in a series of enzymatic, secretory, transcriptional and immunological responses to environmentally-induced stimuli (Binns et al. 2009) to restore an organism’s internal homeostasis and tissue integrity. The KEGG metabolic pathways analysis of the transcripts with the highest fold changes between south coast and west coast populations indicated that gene expression profiles of control specimens kept at 20 °C differed between populations at 158 core metabolic pathways. This number increased to 242 core metabolic pathways at 10 °C. At this temperature, the south coast population had 27 upregulated and 172 downregulated KEGG metabolic pathways that were absent in the 20 °C control group. The majority of upregulated metabolic pathways and their associated genes are involved in various signalling pathway (6 genes), polypeptides synthesis (5 genes), immunue responses (4 genes). Similarly, the downregulated metabolic pathways and their associated genes were linked to various core metabolic pathways (14 genes), RNA transport from nucleus to the cytoplasm (13 genes) and surveillance, detection and degradation of the abnormal mRNAs (10 genes) (APPENDIX A-TABLES A8 and A9)..

## Discussion

In the current study, we used genomic data generated by sequencing pooled DNA from four decapod species, and gene expression data from a single species, to investigate local adaptation, genome-wide variation, and population divergence across a thermal gradient at the boundary between the cool-temperate waters of the south-eastern Atlantic and the warm-temperate waters of the westernmost extent of the south-western Indian Ocean.

Our study shows that southern and western populations of the study species form distinct lineages. Despite episodical dispersal of a moderate number of migrants from the south coast to the west coast that is facilitated by the westward flow of nearshore currents (Branch 1984; Roy et al. 2001; Dufois and Rouault 2012), complete homogenization (or panmixia) is not seen for any of these species. The maintenance of the genetic break in the face of sporadic gene flow highlights the critical role of local adaptation in maintaining population structure. Specifically, reduced fitness of recently-dispersed individuals in habitats to which they are not only maladapted, but likely also outcompeted by a local sister population that occupies a very similar ecological niche, facilitates genomic divergence (Teske et al. 2008; Papadopoulos and Teske 2014). This hypothesis is substantiated by clear differences in the gene expression profiles of numerous biological processes and core metabolic pathways between mudprawn ecotypes at low temperatures (10 °C) that the species’ south coast ecotype does not usually experience within its home range.

### Population structure and divergence time estimates

The finding that all four decapod crustaceans have distinct west- and south coast populations contradicts earlier mtDNA evidence. Previously, phylogenetic splits were found only in *C. kraussi* (Teske et al. 2009) and in *U. africana* (Teske et al. 2006; Teske et al. 2007), whereas *H. orbiculare* (Teske et al. 2014) and *D. brevirostris* (Landschoff and Gouws 2018) were genetically homogeneous. This finding contributes to the growing evidence that mtDNA may overlook population differentiation, and confirms the power of large-scale genomic data to elucidate the evolutionary history of marine animals in unprecedented detail (Carreras et al. 2017; Teske et al. 2019; Coscia et al. 2020). Confidence intervals of divergence time estimates are too wide to conclude that divergence into western and eastern populations occurred at different times. Nonetheless, the relatively old age of the ecotypes (Middle to Late Pleistocene) supports the idea that their divergence could be interpreted as early stage of incipient ecological speciation.

The molecular dating approach used in the present study has so far been applied only to a few taxa, and for that reason requires some clarifications (Leblois et al. 2018; Bourgeois et al. 2019; Druet et al. 2020). While the basic assumptions of Kimura’s diffusion model (e.g., lack of migration and recurrent mutation after divergence) appear simplistic when applied to marine populations, the Bayesian method applied in KimTree2 is robust to moderate violations of these assumptions. Gautier and Vitalis (2012) and Clemente et al. (2018b) estimated correct branch lengths under low to moderate migration rates (*M* = *N*_*e*_*m* between 0.1 and 1), and only very high migration rates (e.g., M = 10) resulted in underestimates in branch lengths. The estimated *F*_ST_ between pairs of populations suggests that migration is low to moderate. Moreover, sampling sites were located well within the cool-temperate and warm-temperate marine biogeographic provinces, respectively, rather than the contact zone on the south-west coast where the overlap in the range of populations facilitates introgression (Teske et al. 2014), which would affect branch length estimates. Violation of neutrality does not negatively impact branch length estimates, provided that only a small proportion of loci are subject to the selection (Gautier et al. 2010), as was also the case here. While local dependency between neighbouring SNPs in the form of linkage disequilibrium coefficient (*r*^2^) (Hill and Robertson 1968) is not addressed directly by the model applied in KimTree2, Gautier and Vitalis (2012) suggested that this will have limited influence on the estimation of the branch length. Moreover, Clemente et al. (2018a) showed that interdependence between adjacent SNPs is more likely to affect the precision of posterior distribution than the accuracy of branch length estimates. Rigorous filtering of SNPs in the current dataset ensured that only sub-selected loci that have been successfully genotyped in both populations also acted in favour of independence of the selected markers.

### Thermal adaptation and incipient ecological speciation

Clear differences in gene expression profiles were identified between the west coast and south coast populations of *Upogebia africana*, particularly at the lower temperature. Specifically, exposure to 10 °C invoked a disruption in the gene expression profile of the south coast population compared to control specimens that were treated at 20 °C. Some of the major metabolic pathways negatively affected at 10 °C were involved in thermoregulatory, immunological and inflammatory responses, and their interruption negatively affects long-term fitness (e.g. reproductive output, growth, and survival) of individuals that have dispersed into thermally challenging habitat to which their genome possesses no prior adaptation (Martin et al. 2008). This finding supports the idea that reduced fitness of migrants from the south coast that have settled on the west coast excludes them from this region during episodes of strong cold-water upwelling typically occurs in this bioregion, and drives the process of incipient speciation.

The functional significance of transcriptomic responses in non-model species without annotated reference genomes largely relies on predicted protein structure and function based on homology to published data on proteins and metabolic pathways from closely related species. Because of this, such results need to be interpreted within these limitations. Nevertheless, some of the genes involved in thermoregulatory pathways demonstrated heterogeneous gene expression responses to the temperature treatments. For instance, in the specimens exposed to10 °C, the products of several genes in the heat shock chaperon protein family (HS90B, HSP70, HSP71, HSP76, HSP7C, HSP7D, HSP83) and an immune system-related gene, with high homology to the vertebrates class II histocompatibility gene (HG2A), were significantly upregulated. While at 20 °C only the product of one of these genes, HSP70, was significantly upregulated. This points to complex transcriptomic responses of this group of proteins, and other similar-acting polypeptides, to environmental stressors, as found in numerous previous studies (Ovelgönne et al. 2000; Mahat et al. 2016; Gierz et al. 2017; Lee et al. 2017).

### Conclusion

Although thermal adaptation has long been assumed to be an important driver of speciation in marine regions that lack absolute dispersal barriers (Teske et al. 2011), distinguishing its heritable nature from phenotypic plasticity has only become possible with recent advancements in massively-parallel sequencing technology (Sandoval-Castillo et al. 2020). It is commonly assumed that movement of one effective migrant per generation is sufficient to homogenize population differentiation that follows genetic drift (Wright 1931; Slatkin 1985). The present study shows that gene flow is not necessarily required to explain the presence of population structure across the boundary between the Atlantic and Indian Oceans that was previously identified using selectively neutral markers (Teske et al., 2011). Instead, it provides clear evidence for the role of post-dispersal selection against migrants that have established themselves in habitat to which they are maladapted.

In addition, the finding that all four crustacean species for which genomic data were generated are genetically structured across this biogeographical boundary contributes to the growing evidence that the application of the data generated through DNA barcoding initiatives for population genetic studies may overlook biologically meaningful population structure (Teske et al. 2019). Importantly, molecular dating shows that Atlantic and Indian Ocean ecotypes of the two species that show no mtDNA-based genetic structure across this boundary (*H. orbiculare* and *D. brevirostris*) did not diverge from each other significantly more recently than the other two species’ ecotypes. These results thus provide a more comprehensive overview of the evolution and maintenance of genetic structure during the early stages of the ecological speciation process across the temperature-defined biogeographical boundary at the southern tip of Africa.

## Data availability

Raw sequences generated for this study were submitted to the NCBI Short Reads Archive (SRA) under Bioporojects PRJNA660321 and PRJNA660316. [These data will be available publicly before the publication]

## Author contributions

P.R.T. conceived the research, obtained the funding, collected the samples and conducted the thermal exposure experiments. I.S.K. and R.J.T. generated the Pool-Seq data. A.E.-K. and P.R.T. generated the RNAseq data. A.E.-K. and D.M.M. analysed the data. All authors contributed to the writing of the manuscript, and read and approved the final version.

## Acknowledgements

This study was funded by the PADI Foundation (grant no. 10981 to P.R.T.) and the University of Johannesburg (URC/FRC grant to P.R.T.). A.E.-K. and D.M.M acknowledge the University of Johannesburg for a Global Excellence and Stature (GES) postdoctoral research and doctorate fellowships and South African National Antarctic Program (SANAP) research grants. Computational platforms for this study were provided by the Centre for High-Performance Computation (CHPC) in Cape Town and the University of Johannesburg IT department. The authors would like to express their gratitude to Robert Kofler, Ronald Vitalis and Fabian Clemente for their advice.

