## Supplementary files for "Genomic divergence and differential gene expression between crustacean ecotypes across a marine thermal gradient"

Table A1 Location, number of specimens in each pool, number of quality-filtered sequences, and the mapping statistics for data generated for the four species in the current study.

| **Species Name** | **Population** | **Number of Individuals** | **Number of Reads Passed Initial Quality Filters** | **Percentage of Mapped Reads** | **Average Quality of Mapped Reads in Phred** |
| --- | --- | --- | --- | --- | --- |
|  | South Coast | 14 | 1,853,844 | 93.97 | 56.22 |
| *Callichirus kraussi* |  |  |  |  |  |
|  | West Coast | 14 | 2,538,165 | 94.07 | 58.22 |
|  | South Coast | 20 | 1,601,061 | 97.9 | 42.5 |
| *Upogebia africana* |  |  |  |  |  |
|  | West Coast | 24 | 2,569,490 | 99.66 | 41.92 |
|  | South Coast | 14 | 2,158,460 | 96.67 | 43.43 |
| *Diogenes brevirostris* |  |  |  |  |  |
|  | West Coast | 21 | 2,029,722 | 97.49 | 43.45 |
|  | South Coast | 13 | 1,657,050 | 93.83 | 57.31 |
| *Hymenosoma orbiculare* |  |  |  |  |  |
|  | West Coast | 13 | 1,560,173 | 91.35 | 57.54 |

Table A2 Number of SNPs generated from pool sequencing of each study species.

| **Species Name** | **Total Number Of SNPs** | **Number of Quality Passed SNPs** | **Number of SNPS Genotyped In Both Populations** |
| --- | --- | --- | --- |
| *Callichirus kraussi* | 6,655 | 6,398 | 1,742 |
| *Upogebia africana* | 3,318 | 2,994 | 1,254 |
| *Diogenes brevirostris* | 3,668 | 3,195 | 1,598 |
| *Hymenosoma orbiculare* | 9,831 | 9,251 | 2,829 |

*Table A3* Fst and the percentage of alleles that show significant changes (α < 0.05) in the frequency between pair of populations.

| **Species Name** | **Fst** | **Percentage of Significant Allele Frequency Changes** |
| --- | --- | --- |
| *Callichirus kraussi* | 0.0615(95%CI 0.0637-0.0594) | 3.235 |
| *Upogebia africana* | 0.0341(95%CI 0.0376-0.0306 ) | 2.428 |
| *Diogenes brevirostris* | 0.0399 (95%CI 0.0427-0.037) | 2.0511 |
| *Hymenosoma orbiculare* | 0.0612(95%CI 0.063589-0.059) | 4.179 |

Table A4 Total number of MCMC iterations, burn-in steps, initial seeds, and Gelman and Rubin's shrinking factors for KimTree2 branch length estimates of each study species.

| **Species Name** | **Population** | **Number of Iteration** | **Burn-in** | **Initial Seed For Three Chains** | **Shrinking Factor-(Upper Lower CI)** | **Multivariate psrf** |
| --- | --- | --- | --- | --- | --- | --- |
|  | South Coast |  |  |  | 1.01 -1.03 |  |
| *Callichirus kraussi* |  | (3x10)^11 | (1x10)^11 | 1562347843_1562615643_server clock |  | 1.01 |
|  | West Coast |  |  |  | 1.01 - 1.02 |  |
|  | South Coast |  |  |  | 1.01 - 1.05 |  |
| *Upogebia africana* |  | (7.5x10)^11 | (2.5x10)^11 | 1561624788_1561624830_1564987633 |  | 1.05 |
|  | West Coast |  |  |  | 1.02 -1.08 |  |
|  | South Coast |  |  |  | 1.03 -1.1 |  |
| *Diogenes brevirostris* |  | (7.5x10)^11 | (2.5x10)^11 | 1563005438_1593693594_1284865476 |  | 1.08 |
|  | West Coast |  |  |  | 1.04 -1.1 |  |
|  | South Coast |  |  |  | 1.01-1.03 |  |
| *Hymenosoma orbiculare* |  | (3x10)^11 | (1x10)^11 | 1561654729_1563691721_1563691698 |  | 1.02 |
|  | West Coast |  |  |  | 1.01-1.03 |  |

Table A5 Branch length estimates$(\boldsymbol{\tau}_{\mathbf{i}})$ for branches leading to southern and westerns population of each study species.

| **Species Name** | **Population** | **Branch length in Diffusion Time Scale (Average over 3 chains)** |
| --- | --- | --- |
|  | South Coast | 0.171 |
| *Callichirus kraussi* |  |  |
|  | West Coast | 0.157 |
|  | South Coast | 0.266 |
| *Upogebia africana* |  |  |
|  | West Coast | 0.258 |
|  | South Coast | 0.3868 |
| *Diogenes brevirostris* |  |  |
|  | West Coast | 0.3872 |
|  | South Coast | 0.185 |
| *Hymenosoma orbiculare* |  |  |
|  | West Coast | 0.169 |

Table A6 Assembly statistics of Upogebia africana hepatopancereas transcriptome.

| **Statistics** | **Value** |
| --- | --- |
| # contigs ( ≥ 0 bp) | 664,707 |
| # contigs ( ≥ 1000 bp) | 75,293 |
| # contigs ( ≥ 5000 bp) | 802 |
| # contigs ( ≥ 10000 bp) | 22 |
| # contigs ( ≥ 25000 bp) | 2 |
| Total length ( ≥ 0 bp) | 368,811,389 |
| Largest contig | 48,515 |
| GC (%) | 44.95 |
| N50 | 1,152 |
| N75 | 732 |
| L50 | 59,198 |
| L75 | 122,365 |

Table A7 Functional annotation of Differentially Expressed (DE) transcripts in terms of Gene Ontology Categorisation, Biological Process, in Upogebia africana hepatopancreas transcriptome treated at different temperature.

| **Biological Process** | **Percentage of DE genes at 10 °C** | **Percentage of DE genes at 20 °C** |
| --- | --- | --- |
| cellular process | 39.4 | 34.1 |
| biological regulation | 15.9 | 12.2 |
| regulation of biological process | 13.9 | 9.3 |
| regulation of biological process | 13.9 | 9.3 |
| response to stimulus | 11.2 | 7.6 |
| regulation of cellular process | 11.2 | 6.8 |
| regulation of cellular process | 11.2 | 6.8 |
| cellular component organization | 9.9 | 7.5 |
| cellular component organization | 9.9 | 7.5 |
| developmental process | 7.7 | 5.0 |
| multicellular organismal process | 7.5 | 5.1 |
| anatomical structure development | 7.0 | 4.4 |
| cellular response to stimulus | 6.9 | 4.4 |
| cellular response to stimulus | 6.9 | 4.4 |
| negative regulation of biological process | 5.3 | 3.1 |
| negative regulation of biological process | 5.3 | 3.1 |
| positive regulation of biological process | 5.3 | 2.7 |
| positive regulation of biological process | 5.3 | 2.7 |
| multicellular organism development | 5.2 | 3.6 |
| response to stress | 5.1 | 3.4 |
| localization | 5.0 | 8.0 |
| response to chemical | 4.3 | 2.8 |
| positive regulation of cellular process | 4.2 | 2.1 |
| positive regulation of cellular process | 4.2 | 2.1 |
| establishment of localization | 4.1 | 6.3 |
| cellular developmental process | 3.6 | 1.8 |
| cellular developmental process | 3.6 | 1.8 |
| negative regulation of cellular process | 3.6 | 1.7 |
| negative regulation of cellular process | 3.6 | 1.7 |
| cell communication | 2.9 | 1.4 |
| positive regulation of metabolic process | 2.7 | 1.3 |
| positive regulation of metabolic process | 2.7 | 1.3 |
| signaling | 2.5 | 1.2 |
| signal transduction | 2.3 | 1.1 |
| signal transduction | 2.3 | 1.1 |
| immune system process | 2.1 | 0.6 |
| regulation of signaling | 1.4 | 0.5 |
| regulation of signaling | 1.4 | 0.5 |
| transporter activity | 1.2 | 3.1 |
| transmembrane transporter activity | 0.9 | 2.8 |
| cellular homeostasis | 0.4 | 1.1 |
| cell division | 0.4 | 1.0 |

Table A8 Functional annotation of Differentially Expressed (DE) transcripts in term of KEGG metabolic pathways and their associated genes in hepatopancreas transcriptome of Upogebia africana treated at 20 °C.

| **Gene Name** |  | **Definition** | **Homology-based functional annotation** |
| --- | --- | --- | --- |
| DIO1 |  | type I thyroxine 5'-deiodinase [EC:1.21.99.4] | Thyroid_hormone_signaling_pathway(3) |
| PGAM, gpmA |  | 2,3-bisphosphoglycerate-dependent phosphoglycerate mutase [EC:5.4.2.11] | Glycolysis_/_Gluconeogenesis(2) Glycine,_serine_and_threonine_metabolism(1) Methane_metabolism(2) Metabolic_pathways(20) Biosynthesis_of_secondary_metabolites(7) Microbial_metabolism_in_diverse_environments(4) Biosynthesis_of_antibiotics(6) Carbon_metabolism(4) Biosynthesis_of_amino_acids(1) Glucagon_signaling_pathway(3) Central_carbon_metabolism_in_cancer(1) |
| CALM |  | calmodulin | Ras_signaling_pathway(1) Rap1_signaling_pathway(1) MAPK_signaling_pathway_-_plant(1) Calcium_signaling_pathway(2) cGMP-PKG_signaling_pathway(3) cAMP_signaling_pathway(1) Phosphatidylinositol_signaling_system(1) Oocyte_meiosis(2) Cellular_senescence(2) Adrenergic_signaling_in_cardiomyocytes(1) Vascular_smooth_muscle_contraction(2) Apelin_signaling_pathway(1) C-type_lectin_receptor_signaling_pathway(2) Plant-pathogen_interaction(2) Circadian_entrainment(1) Long-term_potentiation(2) Neurotrophin_signaling_pathway(1) Dopaminergic_synapse(2) Olfactory_transduction(1) Phototransduction(1) Phototransduction_-_fly(1) Inflammatory_mediator_regulation_of_TRP_channels(1) Insulin_signaling_pathway(1) GnRH_signaling_pathway(1) Estrogen_signaling_pathway(3) Melanogenesis(1) Oxytocin_signaling_pathway(3) Glucagon_signaling_pathway(3) Renin_secretion(3) Aldosterone_synthesis_and_secretion(1) Salivary_secretion(2) Gastric_acid_secretion(1) Alzheimer_disease(2) Amphetamine_addiction(2) Alcoholism(3) Pertussis(1) Tuberculosis(4) Human_cytomegalovirus_infection(2) Kaposi_sarcoma-associated_herpesvirus_infection(2) Human_immunodeficiency_virus_1_infection(2) Pathways_in_cancer(2) Glioma(1) Fluid_shear_stress_and_atherosclerosis(2) |
| RP-L14e, RPL14 |  | large subunit ribosomal protein L14e | Ribosome(19) |
| RP-L21e, RPL21 |  | large subunit ribosomal protein L21e | Ribosome(19) |
| RP-L23e, RPL23 |  | large subunit ribosomal protein L23e | Ribosome(19) |
| RP-L26e, RPL26 |  | large subunit ribosomal protein L26e | Ribosome(19) |
| RP-L27Ae, RPL27A |  | large subunit ribosomal protein L27Ae | Ribosome(19) |
| RP-L44e, RPL44 |  | large subunit ribosomal protein L44e | Ribosome(19) |
| RP-L6e, RPL6 |  | large subunit ribosomal protein L6e | Ribosome(19) |
| RP-L7e, RPL7 |  | large subunit ribosomal protein L7e | Ribosome(19) |
| RP-LP1, RPLP1 |  | large subunit ribosomal protein LP1 | Ribosome(19) |
| RP-S11e, RPS11 |  | small subunit ribosomal protein S11e | Ribosome(19) |
| RP-S15e, RPS15 |  | small subunit ribosomal protein S15e | Ribosome(19) |
| RP-S20e, RPS20 |  | small subunit ribosomal protein S20e | Ribosome(19) |
| RP-S26e, RPS26 |  | small subunit ribosomal protein S26e | Ribosome(19) |
| RP-S7e, RPS7 |  | small subunit ribosomal protein S7e | Ribosome(19) |
| EEF2 |  | elongation factor 2 | AMPK_signaling_pathway(2) Oxytocin_signaling_pathway(3) |
| PPIA |  | peptidyl-prolyl cis-trans isomerase A (cyclophilin A) [EC:5.2.1.8] | Cationic_antimicrobial_peptide_(CAMP)_resistance(1) Necroptosis(2) |
| PLAUR, CD87 |  | plasminogen activator, urokinase receptor | Complement_and_coagulation_cascades(1) Proteoglycans_in_cancer(5) |
| KCNMA1, KCA1.1 |  | potassium large conductance calcium-activated channel subfamily M alpha member 1 | cGMP-PKG_signaling_pathway(3) Vascular_smooth_muscle_contraction(2) Insulin_secretion(1) Renin_secretion(3) Salivary_secretion(2) Pancreatic_secretion(2) |
| COL1A |  | collagen, type I, alpha | PI3K-Akt_signaling_pathway(2) Focal_adhesion(2) ECM-receptor_interaction(1) Platelet_activation(1) Relaxin_signaling_pathway(1) AGE-RAGE_signaling_pathway_in_diabetic_complications(1) Protein_digestion_and_absorption(1) Amoebiasis(2) Human_papillomavirus_infection(3) Proteoglycans_in_cancer(5) |
| CD63, MLA1, TSPAN30 |  | CD63 antigen | Lysosome(2) Proteoglycans_in_cancer(5) |
| DIO3 |  | thyroxine 5-deiodinase [EC:1.21.99.3] | Thyroid_hormone_signaling_pathway(3) |
| CHST13 |  | chondroitin 4-sulfotransferase 13 [EC:2.8.2.5] | Glycosaminoglycan_biosynthesis_-_chondroitin_sulfate_/_dermatan_sulfate(1) |
| LUM |  | lumican | Proteoglycans_in_cancer(5) |
| SLC22A2, OCT2 |  | MFS transporter, OCT family, solute carrier family 22 (organic cation transporter), member 2 | Choline_metabolism_in_cancer(2) |
| SLC22A4_5, OCTN |  | MFS transporter, OCT family, solute carrier family 22 (organic cation transporter), member 4/5 | Choline_metabolism_in_cancer(2) |
| SLC22A7, OAT2 |  | MFS transporter, OCT family, solute carrier family 22 (organic anion transporter), member 7 | Bile_secretion(1) |
| ASTL |  | astacin-like metalloendopeptidase [EC:3.4.-.-] |  |
| K08884 |  | serine/threonine protein kinase, bacterial [EC:2.7.11.1] |  |
| CHST8 |  | carbohydrate 4-sulfotransferase 8 [EC:2.8.2.-] | Various_types_of_N-glycan_biosynthesis(1) Metabolic_pathways(20) |
| DYNLL |  | dynein light chain LC8-type | Vasopressin-regulated_water_reabsorption(1) Salmonella_infection(2) |
| BTBD3_6 |  | BTB/POZ domain-containing protein 3/6 |  |
| ERCC3, XPB |  | DNA excision repair protein ERCC-3 [EC:3.6.4.12] | Basal_transcription_factors(4) Nucleotide_excision_repair(6) |
| RBP |  | reticulocyte-binding protein | Malaria(1) |
| sasG |  | surface protein G | Staphylococcus_aureus_infection(1) |
| MPH1 |  | ATP-dependent DNA helicase MPH1 [EC:3.6.4.12] |  |
| ENDOU, PP11 |  | poly(U)-specific endoribonuclease [EC:3.1.-.-] |  |
| ANXA7_11 |  | annexin A7/11 |  |
| LCE |  | choriolysin L [EC:3.4.24.66] |  |
| thyA, TYMS |  | thymidylate synthase [EC:2.1.1.45] | Pyrimidine_metabolism(4) One_carbon_pool_by_folate(1) Metabolic_pathways(20) Antifolate_resistance(1) |
| mtrA |  | tetrahydromethanopterin S-methyltransferase subunit A [EC:2.1.1.86] | Methane_metabolism(2) Metabolic_pathways(20) Microbial_metabolism_in_diverse_environments(4) Carbon_metabolism(4) |
| SPT |  | serine palmitoyltransferase [EC:2.3.1.50] | Sphingolipid_metabolism(2) Metabolic_pathways(20) Sphingolipid_signaling_pathway(2) Autophagy_-_yeast(2) |
| DHPS, dys |  | deoxyhypusine synthase [EC:2.5.1.46] |  |
| tmk, DTYMK |  | dTMP kinase [EC:2.7.4.9] | Pyrimidine_metabolism(4) Metabolic_pathways(20) |
| UGP2, galU, galF |  | UTP--glucose-1-phosphate uridylyltransferase [EC:2.7.7.9] | Pentose_and_glucuronate_interconversions(1) Galactose_metabolism(1) Starch_and_sucrose_metabolism(2) Amino_sugar_and_nucleotide_sugar_metabolism(4) Metabolic_pathways(20) Biosynthesis_of_secondary_metabolites(7) Biosynthesis_of_antibiotics(6) |
| PGLS, pgl, devB |  | 6-phosphogluconolactonase [EC:3.1.1.31] | Pentose_phosphate_pathway(2) Metabolic_pathways(20) Biosynthesis_of_secondary_metabolites(7) Microbial_metabolism_in_diverse_environments(4) Biosynthesis_of_antibiotics(6) Carbon_metabolism(4) |
| E3.2.1.14 |  | chitinase [EC:3.2.1.14] | Amino_sugar_and_nucleotide_sugar_metabolism(4) Metabolic_pathways(20) |
| map |  | methionyl aminopeptidase [EC:3.4.11.18] |  |
| ppa |  | inorganic pyrophosphatase [EC:3.6.1.1] | Oxidative_phosphorylation(2) |
| dut, DUT |  | dUTP pyrophosphatase [EC:3.6.1.23] | Pyrimidine_metabolism(4) Drug_metabolism_-_other_enzymes(2) Metabolic_pathways(20) |
| E7.6.2.1 |  | phospholipid-translocating ATPase [EC:7.6.2.1] |  |
| ATP2C |  | P-type Ca2+ transporter type 2C [EC:7.2.2.10] |  |
| GPI, pgi |  | glucose-6-phosphate isomerase [EC:5.3.1.9] | Glycolysis_/_Gluconeogenesis(2) Pentose_phosphate_pathway(2) Starch_and_sucrose_metabolism(2) Amino_sugar_and_nucleotide_sugar_metabolism(4) Metabolic_pathways(20) Biosynthesis_of_secondary_metabolites(7) Microbial_metabolism_in_diverse_environments(4) Biosynthesis_of_antibiotics(6) Carbon_metabolism(4) |
| ATPeV0A, ATP6N |  | V-type H+-transporting ATPase subunit a | Oxidative_phosphorylation(2) Metabolic_pathways(20) Lysosome(2) Phagosome(1) Synaptic_vesicle_cycle(1) Collecting_duct_acid_secretion(1) Vibrio_cholerae_infection(2) Epithelial_cell_signaling_in_Helicobacter_pylori_infection(1) Tuberculosis(4) Human_papillomavirus_infection(3) Rheumatoid_arthritis(1) |
| CDC7 |  | cell division control protein 7 [EC:2.7.11.1] | Cell_cycle(2) Cell_cycle_-_yeast(2) Meiosis_-_yeast(2) |
| CLB1 |  | G2/mitotic-specific cyclin 1 | MAPK_signaling_pathway_-_yeast(2) Cell_cycle_-_yeast(2) Meiosis_-_yeast(2) |
| POLD1 |  | DNA polymerase delta subunit 1 [EC:2.7.7.7] | DNA_replication(3) Base_excision_repair(1) Nucleotide_excision_repair(6) Mismatch_repair(2) Homologous_recombination(1) |
| PRI2 |  | DNA primase large subunit | DNA_replication(3) |
| PSMA1 |  | 20S proteasome subunit alpha 6 [EC:3.4.25.1] | Proteasome(8) |
| PSMA4 |  | 20S proteasome subunit alpha 3 [EC:3.4.25.1] | Proteasome(8) |
| PSMB4 |  | 20S proteasome subunit beta 7 [EC:3.4.25.1] | Proteasome(8) |
| PSMB6 |  | 20S proteasome subunit beta 1 [EC:3.4.25.1] | Proteasome(8) |
| RP-L18e, RPL18 |  | large subunit ribosomal protein L18e | Ribosome(19) |
| RP-L27Ae, RPL27A |  | large subunit ribosomal protein L27Ae | Ribosome(19) |
| RP-L44e, RPL44 |  | large subunit ribosomal protein L44e | Ribosome(19) |
| RP-L4e, RPL4 |  | large subunit ribosomal protein L4e | Ribosome(19) |
| RP-S13e, RPS13 |  | small subunit ribosomal protein S13e | Ribosome(19) |
| RPA12, ZNRD1 |  | DNA-directed RNA polymerase I subunit RPA12 | RNA_polymerase(3) |
| RPB7, POLR2G |  | DNA-directed RNA polymerase II subunit RPB7 | RNA_polymerase(3) Huntington_disease(1) |
| RPC6, POLR3F |  | DNA-directed RNA polymerase III subunit RPC6 | RNA_polymerase(3) Cytosolic_DNA-sensing_pathway(1) |
| PSMD14, RPN11, POH1 |  | 26S proteasome regulatory subunit N11 | Proteasome(8) Epstein-Barr_virus_infection(4) |
| PSMD1, RPN2 |  | 26S proteasome regulatory subunit N2 | Proteasome(8) Epstein-Barr_virus_infection(4) |
| PSMC4, RPT3 |  | 26S proteasome regulatory subunit T3 | Proteasome(8) Epstein-Barr_virus_infection(4) |
| PSMC3, RPT5 |  | 26S proteasome regulatory subunit T5 | Proteasome(8) Epstein-Barr_virus_infection(4) |
| TAF5 |  | transcription initiation factor TFIID subunit 5 | Basal_transcription_factors(4) |
| TFIIH2, GTF2H2, SSL1 |  | transcription initiation factor TFIIH subunit 2 | Basal_transcription_factors(4) Nucleotide_excision_repair(6) Viral_carcinogenesis(5) |
| EIF2S3 |  | translation initiation factor 2 subunit 3 | RNA_transport(1) |
| HSPA1s |  | heat shock 70kDa protein 1/2/6/8 | Spliceosome(2) MAPK_signaling_pathway(2) Protein_processing_in_endoplasmic_reticulum(5) Endocytosis(2) Longevity_regulating_pathway_-_multiple_species(1) Antigen_processing_and_presentation(3) Estrogen_signaling_pathway(3) Legionellosis(1) Toxoplasmosis(1) Measles(1) |
| TC.AAA |  | ATP:ADP antiporter, AAA family |  |
| RRP4, EXOSC2 |  | exosome complex component RRP4 | RNA_degradation(4) |
| PIGA, GPI3 |  | phosphatidylinositol N-acetylglucosaminyltransferase subunit A [EC:2.4.1.198] | Glycosylphosphatidylinositol_(GPI)-anchor_biosynthesis(1) Metabolic_pathways(20) |
| HSP90A, htpG |  | molecular chaperone HtpG | Protein_processing_in_endoplasmic_reticulum(5) PI3K-Akt_signaling_pathway(2) Necroptosis(2) Antigen_processing_and_presentation(3) NOD-like_receptor_signaling_pathway(2) Plant-pathogen_interaction(2) IL-17_signaling_pathway(1) Th17_cell_differentiation(2) Progesterone-mediated_oocyte_maturation(1) Estrogen_signaling_pathway(3) Salmonella_infection(2) Pathways_in_cancer(2) Prostate_cancer(1) Fluid_shear_stress_and_atherosclerosis(2) |
| PPP3C, CNA |  | serine/threonine-protein phosphatase 2B catalytic subunit [EC:3.1.3.16] | MAPK_signaling_pathway(2) Calcium_signaling_pathway(2) cGMP-PKG_signaling_pathway(3) Oocyte_meiosis(2) Cellular_senescence(2) Wnt_signaling_pathway(1) Axon_guidance(1) VEGF_signaling_pathway(1) Osteoclast_differentiation(1) C-type_lectin_receptor_signaling_pathway(2) Natural_killer_cell_mediated_cytotoxicity(1) Th1_and_Th2_cell_differentiation(1) Th17_cell_differentiation(2) T_cell_receptor_signaling_pathway(1) B_cell_receptor_signaling_pathway(1) Long-term_potentiation(2) Glutamatergic_synapse(1) Dopaminergic_synapse(2) Oxytocin_signaling_pathway(3) Glucagon_signaling_pathway(3) Renin_secretion(3) Alzheimer_disease(2) Amyotrophic_lateral_sclerosis_(ALS)(1) Amphetamine_addiction(2) Tuberculosis(4) Human_cytomegalovirus_infection(2) Human_T-cell_leukemia_virus_1_infection(2) Kaposi_sarcoma-associated_herpesvirus_infection(2) Human_immunodeficiency_virus_1_infection(2) PD-L1_expression_and_PD-1_checkpoint_pathway_in_cancer(1) |
| CERS1_2_3_4, LASS1_2_3_4 |  | ceramide synthetase [EC:2.3.1.24] | Sphingolipid_metabolism(2) Metabolic_pathways(20) Sphingolipid_signaling_pathway(2) |
| COPA, RET1 |  | coatomer subunit alpha |  |
| DUS1 |  | tRNA-dihydrouridine synthase 1 [EC:1.3.1.88] |  |
| ACTN1_4 |  | actinin alpha 1/4 | Focal_adhesion(2) Adherens_junction(1) Tight_junction(2) Leukocyte_transendothelial_migration(1) Regulation_of_actin_cytoskeleton(2) Shigellosis(4) Amoebiasis(2) Viral_carcinogenesis(5) Systemic_lupus_erythematosus(3) |
| IQGAP2_3 |  | Ras GTPase-activating-like protein IQGAP2/3 | Regulation_of_actin_cytoskeleton(2) |
| PCAF, KAT2, GCN5 |  | histone acetyltransferase [EC:2.3.1.48] | Notch_signaling_pathway(1) Thyroid_hormone_signaling_pathway(3) Human_T-cell_leukemia_virus_1_infection(2) Viral_carcinogenesis(5) |
| truD, PUS7 |  | tRNA pseudouridine13 synthase [EC:5.4.99.27] |  |
| ASK, DBF4 |  | activator of S phase kinase | Cell_cycle(2) |
| K06944 |  | uncharacterized protein |  |
| K07126 |  | uncharacterized protein |  |
| DPH1, dph2 |  | 2-(3-amino-3-carboxypropyl)histidine synthase [EC:2.5.1.108] |  |
| NIP7 |  | 60S ribosome subunit biogenesis protein NIP7 |  |
| K07575 |  | PUA domain protein |  |
| RAB8A, MEL |  | Ras-related protein Rab-8A | Autophagy_-_animal(2) Endocytosis(2) AMPK_signaling_pathway(2) Tight_junction(2) Pancreatic_secretion(2) |
| NFYC |  | nuclear transcription factor Y, gamma | Antigen_processing_and_presentation(3) Tuberculosis(4) |
| AURKX |  | aurora kinase, other [EC:2.7.11.1] |  |
| CCT2 |  | T-complex protein 1 subunit beta |  |
| CCT6 |  | T-complex protein 1 subunit zeta |  |
| DNAJC19 |  | DnaJ homolog subfamily C member 19 |  |
| lolD |  | lipoprotein-releasing system ATP-binding protein [EC:3.6.3.-] | ABC_transporters(1) |
| ACTF |  | actin, other eukaryote |  |
| MYO5 |  | myosin V | Pathogenic_Escherichia_coli_infection(1) |
| TUBG |  | tubulin gamma | Human_papillomavirus_infection(3) |
| KIFC1 |  | kinesin family member C1 |  |
| UBE2A, UBC2, RAD6A |  | ubiquitin-conjugating enzyme E2 A [EC:2.3.2.23] | Ubiquitin_mediated_proteolysis(2) |
| UBE2G1, UBC7 |  | ubiquitin-conjugating enzyme E2 G1 [EC:2.3.2.23] | Ubiquitin_mediated_proteolysis(2) Protein_processing_in_endoplasmic_reticulum(5) Parkinson_disease(1) |
| CNOT4, NOT4, MOT2 |  | CCR4-NOT transcription complex subunit 4 [EC:2.3.2.27] | RNA_degradation(4) |
| RFC2_4 |  | replication factor C subunit 2/4 | DNA_replication(3) Nucleotide_excision_repair(6) Mismatch_repair(2) |
| RRM2 |  | ribonucleoside-diphosphate reductase subunit M2 [EC:1.17.4.1] | Purine_metabolism(2) Pyrimidine_metabolism(4) Glutathione_metabolism(2) Drug_metabolism_-_other_enzymes(2) Metabolic_pathways(20) p53_signaling_pathway(1) |
| ERCC2, XPD |  | DNA excision repair protein ERCC-2 [EC:3.6.4.12] | Basal_transcription_factors(4) Nucleotide_excision_repair(6) |
| XPA |  | DNA-repair protein complementing XP-A cells | Platinum_drug_resistance(1) Nucleotide_excision_repair(6) |
| KDELR |  | ER lumen protein retaining receptor | Vibrio_cholerae_infection(2) |
| LAP3 |  | cytosol aminopeptidase [EC:3.4.11.1 3.4.11.5] | Arginine_and_proline_metabolism(1) Glutathione_metabolism(2) Metabolic_pathways(20) |
| STE12 |  | transcription factor STE12 | MAPK_signaling_pathway_-_yeast(2) |
| H3 |  | histone H3 | Alcoholism(3) Shigellosis(4) Transcriptional_misregulation_in_cancer(2) Systemic_lupus_erythematosus(3) |
| H4 |  | histone H4 | Alcoholism(3) Viral_carcinogenesis(5) Systemic_lupus_erythematosus(3) |
| MYST1, MOF, KAT8 |  | histone acetyltransferase MYST1 [EC:2.3.1.48] |  |
| BRPF3 |  | bromodomain and PHD finger-containing protein 3 |  |
| EZH2 |  | [histone H3]-lysine27 N-trimethyltransferase EZH2 [EC:2.1.1.356] | Lysine_degradation(1) Metabolic_pathways(20) MicroRNAs_in_cancer(1) |
| NUF2, CDCA1 |  | kinetochore protein Nuf2 |  |
| DDX3X, bel |  | ATP-dependent RNA helicase DDX3X [EC:3.6.4.13] | RIG-I-like_receptor_signaling_pathway(1) Hepatitis_B(1) Viral_carcinogenesis(5) |
| INO80, INOC1 |  | DNA helicase INO80 [EC:3.6.4.12] |  |
| USP5_13, UBP14 |  | ubiquitin carboxyl-terminal hydrolase 5/13 [EC:3.4.19.12] |  |
| USP36_42 |  | ubiquitin carboxyl-terminal hydrolase 36/42 [EC:3.4.19.12] |  |
| PNO1, DIM2 |  | RNA-binding protein PNO1 |  |
| ANUBL1 |  | AN1-type zinc finger and ubiquitin domain-containing protein 1 |  |
| SEC62 |  | translocation protein SEC62 | Protein_export(1) Protein_processing_in_endoplasmic_reticulum(5) |
| CNOT3, NOT3 |  | CCR4-NOT transcription complex subunit 3 | RNA_degradation(4) |
| XRN2, RAT1 |  | 5'-3' exoribonuclease 2 [EC:3.1.13.-] | Ribosome_biogenesis_in_eukaryotes(4) RNA_degradation(4) |
| DDX5, DBP2 |  | ATP-dependent RNA helicase DDX5/DBP2 [EC:3.6.4.13] | Spliceosome(2) Transcriptional_misregulation_in_cancer(2) Proteoglycans_in_cancer(5) |
| CPSF3L, INTS11 |  | integrator complex subunit 11 [EC:3.1.27.-] |  |
| WDR77, MEP50 |  | methylosome protein 50 |  |
| LCLAT1, AGPAT8 |  | lysocardiolipin and lysophospholipid acyltransferase [EC:2.3.1.- 2.3.1.51] | Glycerolipid_metabolism(1) Glycerophospholipid_metabolism(1) Metabolic_pathways(20) Biosynthesis_of_secondary_metabolites(7) |
| UFD1 |  | ubiquitin fusion degradation protein 1 | Protein_processing_in_endoplasmic_reticulum(5) |
| PAP |  | poly(A) polymerase [EC:2.7.7.19] | mRNA_surveillance_pathway(2) |
| FIP1L1, FIP1 |  | pre-mRNA 3'-end-processing factor FIP1 | mRNA_surveillance_pathway(2) |
| SMARCAL1, HARP |  | SWI/SNF-related matrix-associated actin-dependent regulator of chromatin subfamily A-like protein 1 [EC:3.6.4.12] |  |
| NOP58 |  | nucleolar protein 58 | Ribosome_biogenesis_in_eukaryotes(4) |
| BMS1 |  | ribosome biogenesis protein BMS1 | Ribosome_biogenesis_in_eukaryotes(4) |
| DDX49, DBP8 |  | ATP-dependent RNA helicase DDX49/DBP8 [EC:3.6.4.13] |  |
| MRD1, RBM19 |  | multiple RNA-binding domain-containing protein 1 |  |
| RPF1 |  | ribosome production factor 1 |  |
| BTAF1, MOT1 |  | TATA-binding protein-associated factor [EC:3.6.4.-] |  |
| SLC35C2 |  | solute carrier family 35, member C2 |  |
| TRM5, TRMT5 |  | tRNA (guanine37-N1)-methyltransferase [EC:2.1.1.228] |  |
| RHA1 |  | RING-H2 zinc finger protein RHA1 |  |
| TUBGCP3, GCP3 |  | gamma-tubulin complex component 3 |  |
| SEPT7, CDC3 |  | septin 7 | Shigellosis(4) |
| CDC11 |  | cell division control protein 11 |  |
| COPG |  | coatomer subunit gamma |  |
| PMM |  | phosphomannomutase [EC:5.4.2.8] | Fructose_and_mannose_metabolism(1) Amino_sugar_and_nucleotide_sugar_metabolism(4) Metabolic_pathways(20) Biosynthesis_of_secondary_metabolites(7) Biosynthesis_of_antibiotics(6) |
| SPN1, IWS1 |  | transcription factor SPN1 |  |
| PPP1R7, SDS22 |  | protein phosphatase 1 regulatory subunit 7 |  |
| ATG16L1 |  | autophagy-related protein 16-1 | Autophagy_-_other(1) Autophagy_-_yeast(2) Autophagy_-_animal(2) NOD-like_receptor_signaling_pathway(2) Shigellosis(4) |
| AK6, FAP7 |  | adenylate kinase [EC:2.7.4.3] | Purine_metabolism(2) Metabolic_pathways(20) Biosynthesis_of_secondary_metabolites(7) Biosynthesis_of_antibiotics(6) Ribosome_biogenesis_in_eukaryotes(4) |

Table A9 Functional annotation of Differentially Expressed (DE) transcripts in term of KEGG metabolic pathways and their associated genes in hepatopancreas transcriptome of Upogebia africana treated at 10 °C.

| **Gene Name** |  | **Definition** | **Homology-based functional annotation** |
| --- | --- | --- | --- |
| CALM |  | calmodulin | Ras_signaling_pathway(1) Rap1_signaling_pathway(1) MAPK_signaling_pathway_-_plant(1) Calcium_signaling_pathway(2) cGMP-PKG_signaling_pathway(3) cAMP_signaling_pathway(1) Phosphatidylinositol_signaling_system(2) Oocyte_meiosis(8) Cellular_senescence(4) Adrenergic_signaling_in_cardiomyocytes(6) Vascular_smooth_muscle_contraction(2) Apelin_signaling_pathway(1) C-type_lectin_receptor_signaling_pathway(2) Plant-pathogen_interaction(2) Circadian_entrainment(1) Long-term_potentiation(2) Neurotrophin_signaling_pathway(1) Dopaminergic_synapse(4) Olfactory_transduction(1) Phototransduction(1) Phototransduction_-_fly(1) Inflammatory_mediator_regulation_of_TRP_channels(1) Insulin_signaling_pathway(2) GnRH_signaling_pathway(1) Estrogen_signaling_pathway(4) Melanogenesis(1) Oxytocin_signaling_pathway(4) Glucagon_signaling_pathway(2) Renin_secretion(3) Aldosterone_synthesis_and_secretion(1) Salivary_secretion(3) Gastric_acid_secretion(1) Alzheimer_disease(3) Amphetamine_addiction(2) Alcoholism(1) Pertussis(2) Tuberculosis(4) Human_cytomegalovirus_infection(2) Kaposi_sarcoma-associated_herpesvirus_infection(2) Human_immunodeficiency_virus_1_infection(3) Pathways_in_cancer(8) Glioma(2) Fluid_shear_stress_and_atherosclerosis(2) |
| RP-L10e, RPL10 |  | large subunit ribosomal protein L10e | Ribosome(17) |
| RP-L26e, RPL26 |  | large subunit ribosomal protein L26e | Ribosome(17) |
| RP-L27Ae, RPL27A |  | large subunit ribosomal protein L27Ae | Ribosome(17) |
| RP-L3e, RPL3 |  | large subunit ribosomal protein L3e | Ribosome(17) |
| RP-L6e, RPL6 |  | large subunit ribosomal protein L6e | Ribosome(17) |
| RP-L8e, RPL8 |  | large subunit ribosomal protein L8e | Ribosome(17) |
| RP-S25e, RPS25 |  | small subunit ribosomal protein S25e | Ribosome(17) |
| RP-S3e, RPS3 |  | small subunit ribosomal protein S3e | Ribosome(17) Pathogenic_Escherichia_coli_infection(5) Salmonella_infection(2) |
| rpoS |  | RNA polymerase nonessential primary-like sigma factor | Biofilm_formation_-_Escherichia_coli(1) Biofilm_formation_-_Vibrio_cholerae(1) |
| EEF1A |  | elongation factor 1-alpha | RNA_transport(13) Legionellosis(2) Leishmaniasis(1) |
| EEF2 |  | elongation factor 2 | AMPK_signaling_pathway(4) Oxytocin_signaling_pathway(4) |
| HSPA1s |  | heat shock 70kDa protein 1/2/6/8 | Spliceosome(4) MAPK_signaling_pathway(3) Protein_processing_in_endoplasmic_reticulum(6) Endocytosis(1) Longevity_regulating_pathway_-_multiple_species(1) Antigen_processing_and_presentation(4) Estrogen_signaling_pathway(4) Legionellosis(2) Toxoplasmosis(1) Measles(3) |
| PLAUR, CD87 |  | plasminogen activator, urokinase receptor | Complement_and_coagulation_cascades(1) Proteoglycans_in_cancer(4) |
| HSP90A, htpG |  | molecular chaperone HtpG | Protein_processing_in_endoplasmic_reticulum(6) PI3K-Akt_signaling_pathway(8) Necroptosis(1) Antigen_processing_and_presentation(4) NOD-like_receptor_signaling_pathway(1) Plant-pathogen_interaction(2) IL-17_signaling_pathway(1) Th17_cell_differentiation(2) Progesterone-mediated_oocyte_maturation(4) Estrogen_signaling_pathway(4) Salmonella_infection(2) Pathways_in_cancer(8) Prostate_cancer(2) Fluid_shear_stress_and_atherosclerosis(2) |
| APOE |  | apolipoprotein E | Cholesterol_metabolism(1) Alzheimer_disease(3) |
| SERPINA |  | serpin A |  |
| KCNMA1, KCA1.1 |  | potassium large conductance calcium-activated channel subfamily M alpha member 1 | cGMP-PKG_signaling_pathway(3) Vascular_smooth_muscle_contraction(2) Insulin_secretion(1) Renin_secretion(3) Salivary_secretion(3) Pancreatic_secretion(1) |
| CFL |  | cofilin | Axon_guidance(2) Fc_gamma_R-mediated_phagocytosis(1) Regulation_of_actin_cytoskeleton(1) Pertussis(2) Human_immunodeficiency_virus_1_infection(3) |
| COL1A |  | collagen, type I, alpha | PI3K-Akt_signaling_pathway(8) Focal_adhesion(3) ECM-receptor_interaction(3) Platelet_activation(1) Relaxin_signaling_pathway(1) AGE-RAGE_signaling_pathway_in_diabetic_complications(1) Protein_digestion_and_absorption(3) Amoebiasis(1) Human_papillomavirus_infection(10) Proteoglycans_in_cancer(4) |
| COL6A |  | collagen, type VI, alpha | PI3K-Akt_signaling_pathway(8) Focal_adhesion(3) ECM-receptor_interaction(3) Protein_digestion_and_absorption(3) Human_papillomavirus_infection(10) |
| CD74, DHLAG |  | CD74 antigen | Antigen_processing_and_presentation(4) Tuberculosis(4) Herpes_simplex_virus_1_infection(2) |
| KRT1 |  | type I keratin, acidic | Estrogen_signaling_pathway(4) Staphylococcus_aureus_infection(2) |
| KRT2 |  | type II keratin, basic |  |
| CHST13 |  | chondroitin 4-sulfotransferase 13 [EC:2.8.2.5] | Glycosaminoglycan_biosynthesis_-_chondroitin_sulfate_/_dermatan_sulfate(1) |
| IFI30, GILT |  | interferon, gamma-inducible protein 30 | Antigen_processing_and_presentation(4) |
| TPM3 |  | tropomyosin 3 | Cardiac_muscle_contraction(3) Adrenergic_signaling_in_cardiomyocytes(6) Pathways_in_cancer(8) Thyroid_cancer(2) Hypertrophic_cardiomyopathy_(HCM)(3) Dilated_cardiomyopathy_(DCM)(3) |
| CCT2 |  | T-complex protein 1 subunit beta |  |
| CHST8 |  | carbohydrate 4-sulfotransferase 8 [EC:2.8.2.-] | Various_types_of_N-glycan_biosynthesis(1) Metabolic_pathways(17) |
| TPM1 |  | tropomyosin 1 | Cardiac_muscle_contraction(3) Adrenergic_signaling_in_cardiomyocytes(6) MicroRNAs_in_cancer(1) Hypertrophic_cardiomyopathy_(HCM)(3) Dilated_cardiomyopathy_(DCM)(3) |
| TPM4 |  | tropomyosin 4 | Cardiac_muscle_contraction(3) Adrenergic_signaling_in_cardiomyocytes(6) Hypertrophic_cardiomyopathy_(HCM)(3) Dilated_cardiomyopathy_(DCM)(3) |
| ERCC3, XPB |  | DNA excision repair protein ERCC-3 [EC:3.6.4.12] | Basal_transcription_factors(11) Nucleotide_excision_repair(8) |
| SLC28A |  | pyrimidine nucleoside transport protein |  |
| HACE1 |  | E3 ubiquitin-protein ligase HACE1 [EC:2.3.2.26] |  |
| CIRBP |  | cold-inducible RNA-binding protein |  |
| PTGR1, LTB4DH |  | prostaglandin reductase 1 [EC:1.3.1.74 1.3.1.48] |  |
| sasG |  | surface protein G | Staphylococcus_aureus_infection(2) |
| MPH1 |  | ATP-dependent DNA helicase MPH1 [EC:3.6.4.12] |  |
| TYW3 |  | tRNA wybutosine-synthesizing protein 3 [EC:2.1.1.282] |  |
| COL2A |  | collagen, type II, alpha | PI3K-Akt_signaling_pathway(8) Focal_adhesion(3) ECM-receptor_interaction(3) Protein_digestion_and_absorption(3) Human_papillomavirus_infection(10) |
| trxB, TRR |  | thioredoxin reductase (NADPH) [EC:1.8.1.9] | Selenocompound_metabolism(2) |
| TRMT1, trm1 |  | tRNA (guanine26-N2/guanine27-N2)-dimethyltransferase [EC:2.1.1.215 2.1.1.216] |  |
| DPH5 |  | diphthine methyl ester synthase [EC:2.1.1.314] |  |
| DHPS, dys |  | deoxyhypusine synthase [EC:2.5.1.46] |  |
| HK |  | hexokinase [EC:2.7.1.1] | Glycolysis_/_Gluconeogenesis(2) Fructose_and_mannose_metabolism(2) Galactose_metabolism(1) Starch_and_sucrose_metabolism(1) Amino_sugar_and_nucleotide_sugar_metabolism(2) Streptomycin_biosynthesis(1) Neomycin,_kanamycin_and_gentamicin_biosynthesis(1) Metabolic_pathways(17) Biosynthesis_of_secondary_metabolites(6) Microbial_metabolism_in_diverse_environments(2) Biosynthesis_of_antibiotics(2) Carbon_metabolism(2) HIF-1_signaling_pathway(1) Insulin_signaling_pathway(2) Type_II_diabetes_mellitus(1) Carbohydrate_digestion_and_absorption(1) Shigellosis(1) Central_carbon_metabolism_in_cancer(1) |
| udk, UCK |  | uridine kinase [EC:2.7.1.48] | Pyrimidine_metabolism(1) Drug_metabolism_-_other_enzymes(1) Metabolic_pathways(17) |
| E2.7.4.8, gmk |  | guanylate kinase [EC:2.7.4.8] | Purine_metabolism(1) Metabolic_pathways(17) |
| GMPP |  | mannose-1-phosphate guanylyltransferase [EC:2.7.7.13] | Fructose_and_mannose_metabolism(2) Amino_sugar_and_nucleotide_sugar_metabolism(2) Metabolic_pathways(17) Biosynthesis_of_secondary_metabolites(6) |
| pgsA, PGS1 |  | CDP-diacylglycerol---glycerol-3-phosphate 3-phosphatidyltransferase [EC:2.7.8.5] | Glycerophospholipid_metabolism(3) Metabolic_pathways(17) |
| gloB, gloC, HAGH |  | hydroxyacylglutathione hydrolase [EC:3.1.2.6] | Pyruvate_metabolism(1) Metabolic_pathways(17) |
| E3.1.3.48 |  | protein-tyrosine phosphatase [EC:3.1.3.48] | Two-component_system(1) |
| E3.1.3.56 |  | inositol-1,4,5-trisphosphate 5-phosphatase [EC:3.1.3.56] | Inositol_phosphate_metabolism(1) Metabolic_pathways(17) Phosphatidylinositol_signaling_system(2) |
| nfo |  | deoxyribonuclease IV [EC:3.1.21.2] | Base_excision_repair(5) |
| WARS, trpS |  | tryptophanyl-tRNA synthetase [EC:6.1.1.2] | Aminoacyl-tRNA_biosynthesis(7) |
| AARS, alaS |  | alanyl-tRNA synthetase [EC:6.1.1.7] | Aminoacyl-tRNA_biosynthesis(7) |
| VARS, valS |  | valyl-tRNA synthetase [EC:6.1.1.9] | Aminoacyl-tRNA_biosynthesis(7) |
| MARS, metG |  | methionyl-tRNA synthetase [EC:6.1.1.10] | Selenocompound_metabolism(2) Aminoacyl-tRNA_biosynthesis(7) Metabolic_pathways(17) |
| CARS, cysS |  | cysteinyl-tRNA synthetase [EC:6.1.1.16] | Aminoacyl-tRNA_biosynthesis(7) |
| FARSB, pheT |  | phenylalanyl-tRNA synthetase beta chain [EC:6.1.1.20] | Aminoacyl-tRNA_biosynthesis(7) |
| NARS, asnS |  | asparaginyl-tRNA synthetase [EC:6.1.1.22] | Aminoacyl-tRNA_biosynthesis(7) |
| ATPeV1A, ATP6A |  | V-type H+-transporting ATPase subunit A [EC:7.1.2.2] | Oxidative_phosphorylation(3) Metabolic_pathways(17) Phagosome(6) mTOR_signaling_pathway(3) Synaptic_vesicle_cycle(3) Collecting_duct_acid_secretion(3) Vibrio_cholerae_infection(4) Epithelial_cell_signaling_in_Helicobacter_pylori_infection(3) Human_papillomavirus_infection(10) Rheumatoid_arthritis(3) |
| ATPeV1B, ATP6B |  | V-type H+-transporting ATPase subunit B | Oxidative_phosphorylation(3) Metabolic_pathways(17) Phagosome(6) mTOR_signaling_pathway(3) Synaptic_vesicle_cycle(3) Collecting_duct_acid_secretion(3) Vibrio_cholerae_infection(4) Epithelial_cell_signaling_in_Helicobacter_pylori_infection(3) Human_papillomavirus_infection(10) Rheumatoid_arthritis(3) |
| ATPeV1D, ATP6M |  | V-type H+-transporting ATPase subunit D | Oxidative_phosphorylation(3) Metabolic_pathways(17) Phagosome(6) mTOR_signaling_pathway(3) Synaptic_vesicle_cycle(3) Collecting_duct_acid_secretion(3) Vibrio_cholerae_infection(4) Epithelial_cell_signaling_in_Helicobacter_pylori_infection(3) Human_papillomavirus_infection(10) Rheumatoid_arthritis(3) |
| CDK7 |  | cyclin-dependent kinase 7 [EC:2.7.11.22 2.7.11.23] | Basal_transcription_factors(11) Nucleotide_excision_repair(8) Cell_cycle(10) |
| CDK2 |  | cyclin-dependent kinase 2 [EC:2.7.11.22] | FoxO_signaling_pathway(2) Cell_cycle(10) Oocyte_meiosis(8) p53_signaling_pathway(1) PI3K-Akt_signaling_pathway(8) Cellular_senescence(4) Progesterone-mediated_oocyte_maturation(4) Cushing_syndrome(1) Hepatitis_C(5) Hepatitis_B(3) Measles(3) Human_papillomavirus_infection(10) Human_T-cell_leukemia_virus_1_infection(5) Epstein-Barr_virus_infection(7) Pathways_in_cancer(8) Viral_carcinogenesis(7) Prostate_cancer(2) Small_cell_lung_cancer(2) Gastric_cancer(3) |
| CDC6 |  | cell division control protein 6 | Cell_cycle(10) Cell_cycle_-_yeast(8) Meiosis_-_yeast(6) |
| CDC7 |  | cell division control protein 7 [EC:2.7.11.1] | Cell_cycle(10) Cell_cycle_-_yeast(8) Meiosis_-_yeast(6) |
| POLD2 |  | DNA polymerase delta subunit 2 | DNA_replication(8) Base_excision_repair(5) Nucleotide_excision_repair(8) Mismatch_repair(8) Homologous_recombination(5) |
| ESP1 |  | separase [EC:3.4.22.49] | Cell_cycle(10) Cell_cycle_-_yeast(8) Meiosis_-_yeast(6) Oocyte_meiosis(8) Human_T-cell_leukemia_virus_1_infection(5) |
| MCM3 |  | DNA replication licensing factor MCM3 [EC:3.6.4.12] | DNA_replication(8) Cell_cycle(10) Cell_cycle_-_yeast(8) Meiosis_-_yeast(6) |
| PRI1 |  | DNA primase small subunit [EC:2.7.7.102] | DNA_replication(8) |
| PSMA1 |  | 20S proteasome subunit alpha 6 [EC:3.4.25.1] | Proteasome(12) |
| PSMA3 |  | 20S proteasome subunit alpha 7 [EC:3.4.25.1] | Proteasome(12) |
| PSMA4 |  | 20S proteasome subunit alpha 3 [EC:3.4.25.1] | Proteasome(12) |
| PSMA5 |  | 20S proteasome subunit alpha 5 [EC:3.4.25.1] | Proteasome(12) |
| PSMB1 |  | 20S proteasome subunit beta 6 [EC:3.4.25.1] | Proteasome(12) |
| PSMB7 |  | 20S proteasome subunit beta 2 [EC:3.4.25.1] | Proteasome(12) |
| RP-L21e, RPL21 |  | large subunit ribosomal protein L21e | Ribosome(17) |
| RP-L23, MRPL23, rplW |  | large subunit ribosomal protein L23 | Ribosome(17) |
| RP-L23e, RPL23 |  | large subunit ribosomal protein L23e | Ribosome(17) |
| RP-L24e, RPL24 |  | large subunit ribosomal protein L24e | Ribosome(17) |
| RP-L30e, RPL30 |  | large subunit ribosomal protein L30e | Ribosome(17) |
| RP-L4e, RPL4 |  | large subunit ribosomal protein L4e | Ribosome(17) |
| RP-L7e, RPL7 |  | large subunit ribosomal protein L7e | Ribosome(17) |
| RP-S13e, RPS13 |  | small subunit ribosomal protein S13e | Ribosome(17) |
| RP-S17e, RPS17 |  | small subunit ribosomal protein S17e | Ribosome(17) |
| RPB1, POLR2A |  | DNA-directed RNA polymerase II subunit RPB1 [EC:2.7.7.6] | RNA_polymerase(6) Huntington_disease(6) |
| RPB2, POLR2B |  | DNA-directed RNA polymerase II subunit RPB2 [EC:2.7.7.6] | RNA_polymerase(6) Huntington_disease(6) |
| RPB5, POLR2E |  | DNA-directed RNA polymerases I, II, and III subunit RPABC1 | RNA_polymerase(6) Cytosolic_DNA-sensing_pathway(3) Huntington_disease(6) |
| RPB9, POLR2I |  | DNA-directed RNA polymerase II subunit RPB9 | RNA_polymerase(6) Huntington_disease(6) |
| RPC1, POLR3A |  | DNA-directed RNA polymerase III subunit RPC1 [EC:2.7.7.6] | RNA_polymerase(6) Cytosolic_DNA-sensing_pathway(3) |
| RPC2, POLR3B |  | DNA-directed RNA polymerase III subunit RPC2 [EC:2.7.7.6] | RNA_polymerase(6) Cytosolic_DNA-sensing_pathway(3) |
| PSMD14, RPN11, POH1 |  | 26S proteasome regulatory subunit N11 | Proteasome(12) Epstein-Barr_virus_infection(7) |
| PSMC1, RPT2 |  | 26S proteasome regulatory subunit T2 | Proteasome(12) Human_papillomavirus_infection(10) Epstein-Barr_virus_infection(7) Viral_carcinogenesis(7) |
| PSMC4, RPT3 |  | 26S proteasome regulatory subunit T3 | Proteasome(12) Epstein-Barr_virus_infection(7) |
| PSMC6, RPT4 |  | 26S proteasome regulatory subunit T4 | Proteasome(12) Epstein-Barr_virus_infection(7) |
| PSMC5, RPT6 |  | 26S proteasome regulatory subunit T6 | Proteasome(12) Epstein-Barr_virus_infection(7) |
| TFIIA2, GTF2A2, TOA2 |  | transcription initiation factor TFIIA small subunit | Basal_transcription_factors(11) Viral_carcinogenesis(7) |
| TAF1 |  | transcription initiation factor TFIID subunit 1 [EC:2.3.1.48 2.7.11.1] | Basal_transcription_factors(11) |
| TAF2 |  | transcription initiation factor TFIID subunit 2 | Basal_transcription_factors(11) |
| TAF5 |  | transcription initiation factor TFIID subunit 5 | Basal_transcription_factors(11) |
| TAF6 |  | transcription initiation factor TFIID subunit 6 | Basal_transcription_factors(11) |
| TAF10 |  | transcription initiation factor TFIID subunit 10 | Basal_transcription_factors(11) |
| TFIIE1, GTF2E1, TFA1, tfe |  | transcription initiation factor TFIIE subunit alpha | Basal_transcription_factors(11) Viral_carcinogenesis(7) |
| TFIIE2, GTF2E2, TFA2 |  | transcription initiation factor TFIIE subunit beta | Basal_transcription_factors(11) Viral_carcinogenesis(7) |
| TFIIF2, GTF2F2, TFG2 |  | transcription initiation factor TFIIF subunit beta [EC:3.6.4.12] | Basal_transcription_factors(11) |
| TOP2 |  | DNA topoisomerase II [EC:5.6.2.2] | Platinum_drug_resistance(4) |
| UBE1, UBA1 |  | ubiquitin-activating enzyme E1 [EC:6.2.1.45] | Ubiquitin_mediated_proteolysis(5) Parkinson_disease(1) |
| EEF2 |  | elongation factor 2 | AMPK_signaling_pathway(4) Oxytocin_signaling_pathway(4) |
| EIF2S1 |  | translation initiation factor 2 subunit 1 | RNA_transport(13) Autophagy_-_yeast(2) Autophagy_-_animal(2) Protein_processing_in_endoplasmic_reticulum(6) Apoptosis(2) Non-alcoholic_fatty_liver_disease_(NAFLD)(1) Hepatitis_C(5) Measles(3) Influenza_A(3) Herpes_simplex_virus_1_infection(2) |
| EIF2S2 |  | translation initiation factor 2 subunit 2 | RNA_transport(13) |
| EIF2S3 |  | translation initiation factor 2 subunit 3 | RNA_transport(13) |
| EIF5B |  | translation initiation factor 5B | RNA_transport(13) |
| EIF3I |  | translation initiation factor 3 subunit I | RNA_transport(13) |
| EIF4A |  | translation initiation factor 4A | RNA_transport(13) |
| APC10, DOC1 |  | anaphase-promoting complex subunit 10 | Cell_cycle(10) Cell_cycle_-_yeast(8) Meiosis_-_yeast(6) Oocyte_meiosis(8) Ubiquitin_mediated_proteolysis(5) Progesterone-mediated_oocyte_maturation(4) Human_T-cell_leukemia_virus_1_infection(5) |
| psmA, prcA |  | proteasome alpha subunit [EC:3.4.25.1] | Proteasome(12) |
| POLK |  | DNA polymerase kappa [EC:2.7.7.7] | Fanconi_anemia_pathway(8) Epstein-Barr_virus_infection(7) Pathways_in_cancer(8) Transcriptional_misregulation_in_cancer(2) Colorectal_cancer(3) Pancreatic_cancer(2) Endometrial_cancer(2) Glioma(2) Thyroid_cancer(2) Basal_cell_carcinoma(1) Melanoma(1) Chronic_myeloid_leukemia(1) Small_cell_lung_cancer(2) Non-small_cell_lung_cancer(1) Breast_cancer(1) Hepatocellular_carcinoma(1) Gastric_cancer(3) |
| smc |  | chromosome segregation protein |  |
| mutL |  | DNA mismatch repair protein MutL | Mismatch_repair(8) |
| OGG1 |  | N-glycosylase/DNA lyase [EC:3.2.2.- 4.2.99.18] | Base_excision_repair(5) |
| RRP4, EXOSC2 |  | exosome complex component RRP4 | RNA_degradation(7) |
| RRP40, EXOSC3 |  | exosome complex component RRP40 | RNA_degradation(7) |
| rnc, DROSHA, RNT1 |  | ribonuclease III [EC:3.1.26.3] | Ribosome_biogenesis_in_eukaryotes(10) Proteoglycans_in_cancer(4) |
| PIGA, GPI3 |  | phosphatidylinositol N-acetylglucosaminyltransferase subunit A [EC:2.4.1.198] | Glycosylphosphatidylinositol_(GPI)-anchor_biosynthesis(1) Metabolic_pathways(17) |
| dnaK, HSPA9 |  | molecular chaperone DnaK | RNA_degradation(7) Longevity_regulating_pathway_-_worm(2) Tuberculosis(4) |
| PPP3C, CNA |  | serine/threonine-protein phosphatase 2B catalytic subunit [EC:3.1.3.16] | MAPK_signaling_pathway(3) Calcium_signaling_pathway(2) cGMP-PKG_signaling_pathway(3) Oocyte_meiosis(8) Cellular_senescence(4) Wnt_signaling_pathway(3) Axon_guidance(2) VEGF_signaling_pathway(1) Osteoclast_differentiation(1) C-type_lectin_receptor_signaling_pathway(2) Natural_killer_cell_mediated_cytotoxicity(1) Th1_and_Th2_cell_differentiation(1) Th17_cell_differentiation(2) T_cell_receptor_signaling_pathway(1) B_cell_receptor_signaling_pathway(1) Long-term_potentiation(2) Glutamatergic_synapse(1) Dopaminergic_synapse(4) Oxytocin_signaling_pathway(4) Glucagon_signaling_pathway(2) Renin_secretion(3) Alzheimer_disease(3) Amyotrophic_lateral_sclerosis_(ALS)(1) Amphetamine_addiction(2) Tuberculosis(4) Human_cytomegalovirus_infection(2) Human_T-cell_leukemia_virus_1_infection(5) Kaposi_sarcoma-associated_herpesvirus_infection(2) Human_immunodeficiency_virus_1_infection(3) PD-L1_expression_and_PD-1_checkpoint_pathway_in_cancer(1) |
| PPP2R2 |  | serine/threonine-protein phosphatase 2A regulatory subunit B | mRNA_surveillance_pathway(12) Sphingolipid_signaling_pathway(2) Cell_cycle_-_yeast(8) PI3K-Akt_signaling_pathway(8) AMPK_signaling_pathway(4) Adrenergic_signaling_in_cardiomyocytes(6) Hippo_signaling_pathway(3) Hippo_signaling_pathway_-_fly(3) Tight_junction(3) Dopaminergic_synapse(4) Chagas_disease_(American_trypanosomiasis)(2) Hepatitis_C(5) Human_papillomavirus_infection(10) |
| PPP2C |  | serine/threonine-protein phosphatase 2A catalytic subunit [EC:3.1.3.16] | mRNA_surveillance_pathway(12) MAPK_signaling_pathway_-_fly(3) Sphingolipid_signaling_pathway(2) Cell_cycle_-_yeast(8) Meiosis_-_yeast(6) Oocyte_meiosis(8) Autophagy_-_other(1) Autophagy_-_yeast(2) Autophagy_-_animal(2) PI3K-Akt_signaling_pathway(8) AMPK_signaling_pathway(4) Adrenergic_signaling_in_cardiomyocytes(6) TGF-beta_signaling_pathway(1) Hippo_signaling_pathway(3) Hippo_signaling_pathway_-_fly(3) Tight_junction(3) Dopaminergic_synapse(4) Long-term_depression(1) Chagas_disease_(American_trypanosomiasis)(2) Hepatitis_C(5) Human_papillomavirus_infection(10) |
| RAD51 |  | DNA repair protein RAD51 | Homologous_recombination(5) Fanconi_anemia_pathway(8) Pathways_in_cancer(8) Pancreatic_cancer(2) |
| CHD8, HELSNF1 |  | chromodomain helicase DNA binding protein 8 [EC:3.6.4.12] | Wnt_signaling_pathway(3) |
| RUVBL1, RVB1, INO80H |  | RuvB-like protein 1 (pontin 52) | Wnt_signaling_pathway(3) |
| FEN1, RAD2 |  | flap endonuclease-1 [EC:3.-.-.-] | DNA_replication(8) Base_excision_repair(5) Non-homologous_end-joining(1) |
| ABCG2, CD338 |  | ATP-binding cassette, subfamily G (WHITE), member 2 | Antifolate_resistance(1) ABC_transporters(1) Bile_secretion(1) |
| ABCB-BAC |  | ATP-binding cassette, subfamily B, bacterial |  |
| ABCE1, Rli1 |  | ATP-binding cassette, sub-family E, member 1 |  |
| ASK, DBF4 |  | activator of S phase kinase | Cell_cycle(10) |
| PLK1 |  | polo-like kinase 1 [EC:2.7.11.21] | FoxO_signaling_pathway(2) Cell_cycle(10) Oocyte_meiosis(8) Progesterone-mediated_oocyte_maturation(4) |
| SMC2 |  | structural maintenance of chromosome 2 | Cell_cycle_-_yeast(8) |
| ychF |  | ribosome-binding ATPase |  |
| K06944 |  | uncharacterized protein |  |
| PELO, DOM34, pelA |  | protein pelota | mRNA_surveillance_pathway(12) |
| TUBA |  | tubulin alpha | Phagosome(6) Apoptosis(2) Tight_junction(3) Gap_junction(2) Huntington_disease(6) Pathogenic_Escherichia_coli_infection(5) |
| TUBB |  | tubulin beta | Phagosome(6) Gap_junction(2) Huntington_disease(6) Pathogenic_Escherichia_coli_infection(5) |
| RFA1, RPA1, rpa |  | replication factor A1 | DNA_replication(8) Nucleotide_excision_repair(8) Mismatch_repair(8) Homologous_recombination(5) Fanconi_anemia_pathway(8) |
| K07497 |  | putative transposase |  |
| NMD3 |  | nonsense-mediated mRNA decay protein 3 | Ribosome_biogenesis_in_eukaryotes(10) RNA_transport(13) |
| K07575 |  | PUA domain protein |  |
| MLH1 |  | DNA mismatch repair protein MLH1 | Platinum_drug_resistance(4) Mismatch_repair(8) Fanconi_anemia_pathway(8) Pathways_in_cancer(8) Colorectal_cancer(3) Endometrial_cancer(2) Gastric_cancer(3) |
| MSH2 |  | DNA mismatch repair protein MSH2 | Platinum_drug_resistance(4) Mismatch_repair(8) Pathways_in_cancer(8) Colorectal_cancer(3) |
| MSH4 |  | DNA mismatch repair protein MSH4 |  |
| MUS81 |  | crossover junction endonuclease MUS81 [EC:3.1.22.-] | Homologous_recombination(5) Fanconi_anemia_pathway(8) |
| CCT1, TCP1 |  | T-complex protein 1 subunit alpha |  |
| CCT3, TRIC5 |  | T-complex protein 1 subunit gamma |  |
| CCT5 |  | T-complex protein 1 subunit epsilon |  |
| CCT7 |  | T-complex protein 1 subunit eta |  |
| MYO5 |  | myosin V | Pathogenic_Escherichia_coli_infection(5) |
| MAPRE |  | microtubule-associated protein, RP/EB family |  |
| UBE2A, UBC2, RAD6A |  | ubiquitin-conjugating enzyme E2 A [EC:2.3.2.23] | Ubiquitin_mediated_proteolysis(5) |
| CNOT4, NOT4, MOT2 |  | CCR4-NOT transcription complex subunit 4 [EC:2.3.2.27] | RNA_degradation(7) |
| MARCH1_8 |  | E3 ubiquitin-protein ligase MARCH1/8 [EC:2.3.2.27] |  |
| UBLE1B, SAE2, UBA2 |  | ubiquitin-like 1-activating enzyme E1 B [EC:6.2.1.45] | Ubiquitin_mediated_proteolysis(5) |
| UBE1C, UBA3 |  | ubiquitin-activating enzyme E1 C [EC:6.2.1.45] | Ubiquitin_mediated_proteolysis(5) |
| BRE1 |  | E3 ubiquitin-protein ligase BRE1 [EC:2.3.2.27] |  |
| RNASEH2A |  | ribonuclease H2 subunit A [EC:3.1.26.4] | DNA_replication(8) |
| EXO1 |  | exonuclease 1 [EC:3.1.-.-] | Mismatch_repair(8) |
| ASF1 |  | histone chaperone ASF1 |  |
| RFC1 |  | replication factor C subunit 1 | DNA_replication(8) Nucleotide_excision_repair(8) Mismatch_repair(8) |
| RFC3_5 |  | replication factor C subunit 3/5 | DNA_replication(8) Nucleotide_excision_repair(8) Mismatch_repair(8) |
| APEX2 |  | AP endonuclease 2 [EC:4.2.99.18] | Base_excision_repair(5) |
| ERCC4, XPF |  | DNA excision repair protein ERCC-4 [EC:3.1.-.-] | Nucleotide_excision_repair(8) Fanconi_anemia_pathway(8) |
| ERCC1 |  | DNA excision repair protein ERCC-1 | Platinum_drug_resistance(4) Nucleotide_excision_repair(8) Fanconi_anemia_pathway(8) |
| SEC61A |  | protein transport protein SEC61 subunit alpha | Protein_export(3) Protein_processing_in_endoplasmic_reticulum(6) Phagosome(6) Vibrio_cholerae_infection(4) |
| hlyIII |  | hemolysin III |  |
| SMG6, EST1A |  | protein SMG6 [EC:3.1.-.-] | mRNA_surveillance_pathway(12) |
| DKC1, NOLA4, CBF5 |  | H/ACA ribonucleoprotein complex subunit 4 [EC:5.4.99.-] | Ribosome_biogenesis_in_eukaryotes(10) |
| HIRA, HIR1 |  | protein HIRA/HIR1 |  |
| TADA2A, ADA2 |  | transcriptional adapter 2-alpha |  |
| ING4 |  | inhibitor of growth protein 4 |  |
| BRD1, BRPF2 |  | bromodomain-containing protein 1 |  |
| NDC80, HEC1, TID3 |  | kinetochore protein NDC80 |  |
| CBX5, HP1A |  | chromobox protein 5 |  |
| DDX3X, bel |  | ATP-dependent RNA helicase DDX3X [EC:3.6.4.13] | RIG-I-like_receptor_signaling_pathway(1) Hepatitis_B(3) Viral_carcinogenesis(7) |
| SMARCA5, SNF2H, ISWI |  | SWI/SNF-related matrix-associated actin-dependent regulator of chromatin subfamily A member 5 [EC:3.6.4.-] |  |
| DHDDS, RER2, SRT1 |  | ditrans,polycis-polyprenyl diphosphate synthase [EC:2.5.1.87] | Terpenoid_backbone_biosynthesis(1) Biosynthesis_of_secondary_metabolites(6) |
| CNOT7_8, CAF1, POP2 |  | CCR4-NOT transcription complex subunit 7/8 | RNA_degradation(7) |
| CNOT1, NOT1 |  | CCR4-NOT transcription complex subunit 1 | RNA_degradation(7) |
| DHX16 |  | pre-mRNA-splicing factor ATP-dependent RNA helicase DHX16 [EC:3.6.4.13] | Spliceosome(4) |
| DDX5, DBP2 |  | ATP-dependent RNA helicase DDX5/DBP2 [EC:3.6.4.13] | Spliceosome(4) Transcriptional_misregulation_in_cancer(2) Proteoglycans_in_cancer(4) |
| EIF4A3, FAL1 |  | ATP-dependent RNA helicase [EC:3.6.4.13] | RNA_transport(13) mRNA_surveillance_pathway(12) Spliceosome(4) |
| PABPC |  | polyadenylate-binding protein | RNA_transport(13) mRNA_surveillance_pathway(12) RNA_degradation(7) |
| CPSF3L, INTS11 |  | integrator complex subunit 11 [EC:3.1.27.-] |  |
| SEC11, sipW |  | signal peptidase I [EC:3.4.21.89] | Protein_export(3) |
| SRPR |  | signal recognition particle receptor subunit alpha | Protein_export(3) |
| GPAT3_4, AGPAT9, AGPAT6 |  | glycerol-3-phosphate O-acyltransferase 3/4 [EC:2.3.1.15] | Glycerolipid_metabolism(2) Glycerophospholipid_metabolism(3) Metabolic_pathways(17) Biosynthesis_of_secondary_metabolites(6) |
| LCLAT1, AGPAT8 |  | lysocardiolipin and lysophospholipid acyltransferase [EC:2.3.1.- 2.3.1.51] | Glycerolipid_metabolism(2) Glycerophospholipid_metabolism(3) Metabolic_pathways(17) Biosynthesis_of_secondary_metabolites(6) |
| BEST2, VMD2L1 |  | bestrophin-2 | Salivary_secretion(3) |
| DERL2_3 |  | Derlin-2/3 | Protein_processing_in_endoplasmic_reticulum(6) |
| SEC24 |  | protein transport protein SEC24 | Protein_processing_in_endoplasmic_reticulum(6) Pathogenic_Escherichia_coli_infection(5) |
| CTU1, NCS6 |  | cytoplasmic tRNA 2-thiolation protein 1 [EC:2.7.7.-] | Sulfur_relay_system(1) |
| DIM1 |  | 18S rRNA (adenine1779-N6/adenine1780-N6)-dimethyltransferase [EC:2.1.1.183] |  |
| XPO1, CRM1 |  | exportin-1 | Ribosome_biogenesis_in_eukaryotes(10) RNA_transport(13) MAPK_signaling_pathway_-_fly(3) Influenza_A(3) Human_T-cell_leukemia_virus_1_infection(5) |
| NUP98, ADAR2, NUP116 |  | nuclear pore complex protein Nup98-Nup96 | RNA_transport(13) Influenza_A(3) |
| UPF1, RENT1 |  | regulator of nonsense transcripts 1 [EC:3.6.4.-] | RNA_transport(13) mRNA_surveillance_pathway(12) |
| PAP |  | poly(A) polymerase [EC:2.7.7.19] | mRNA_surveillance_pathway(12) |
| CPSF2, CFT2 |  | cleavage and polyadenylation specificity factor subunit 2 | mRNA_surveillance_pathway(12) |
| CPSF3, YSH1 |  | cleavage and polyadenylation specificity factor subunit 3 [EC:3.1.27.-] | mRNA_surveillance_pathway(12) |
| FIP1L1, FIP1 |  | pre-mRNA 3'-end-processing factor FIP1 | mRNA_surveillance_pathway(12) |
| CSTF2, RNA15 |  | cleavage stimulation factor subunit 2 | mRNA_surveillance_pathway(12) |
| UTP21, WDR36 |  | U3 small nucleolar RNA-associated protein 21 | Ribosome_biogenesis_in_eukaryotes(10) |
| DIP2, UTP12, WDR3 |  | U3 small nucleolar RNA-associated protein 12 | Ribosome_biogenesis_in_eukaryotes(10) |
| IMP4 |  | U3 small nucleolar ribonucleoprotein protein IMP4 | Ribosome_biogenesis_in_eukaryotes(10) |
| NOP1, FBL |  | rRNA 2'-O-methyltransferase fibrillarin [EC:2.1.1.-] | Ribosome_biogenesis_in_eukaryotes(10) |
| UTP24, FCF1 |  | U3 small nucleolar RNA-associated protein 24 | Ribosome_biogenesis_in_eukaryotes(10) |
| BMS1 |  | ribosome biogenesis protein BMS1 | Ribosome_biogenesis_in_eukaryotes(10) |
| RIB2, PUS8 |  | tRNA pseudouridine32 synthase / 2,5-diamino-6-(5-phospho-D-ribitylamino)-pyrimidin-4(3H)-one deaminase [EC:5.4.99.28] | Riboflavin_metabolism(1) Metabolic_pathways(17) |
| SLC39A1_2_3, ZIP1_2_3 |  | solute carrier family 39 (zinc transporter), member 1/2/3 |  |
| DDX49, DBP8 |  | ATP-dependent RNA helicase DDX49/DBP8 [EC:3.6.4.13] |  |
| BRX1, BRIX1 |  | ribosome biogenesis protein BRX1 |  |
| BUD20 |  | bud site selection protein 20 |  |
| NSA2 |  | ribosome biogenesis protein NSA2 |  |
| PES1, NOP7 |  | pescadillo |  |
| RSA4, NLE1 |  | ribosome assembly protein 4 |  |
| SPB1, FTSJ3 |  | AdoMet-dependent rRNA methyltransferase SPB1 [EC:2.1.1.-] |  |
| SUPT5H, SPT5 |  | transcription elongation factor SPT5 |  |
| BTAF1, MOT1 |  | TATA-binding protein-associated factor [EC:3.6.4.-] |  |
| BRF1, GTF3B |  | transcription factor IIIB 90 kDa subunit |  |
| BDP1, TFC5 |  | transcription factor TFIIIB component B'' |  |
| BRIP1, BACH1, FANCJ |  | fanconi anemia group J protein [EC:3.6.4.12] | Homologous_recombination(5) Fanconi_anemia_pathway(8) |
| TAD2, ADAT2 |  | tRNA-specific adenosine deaminase 2 [EC:3.5.4.-] |  |
| gpmI |  | 2,3-bisphosphoglycerate-independent phosphoglycerate mutase [EC:5.4.2.12] | Glycolysis_/_Gluconeogenesis(2) Glycine,_serine_and_threonine_metabolism(1) Methane_metabolism(1) Metabolic_pathways(17) Biosynthesis_of_secondary_metabolites(6) Microbial_metabolism_in_diverse_environments(2) Biosynthesis_of_antibiotics(2) Carbon_metabolism(2) Biosynthesis_of_amino_acids(1) |
| RNF121 |  | RING finger protein 121 |  |
| YWHAB_Q_Z |  | 14-3-3 protein beta/theta/zeta | MAPK_signaling_pathway_-_fly(3) Cell_cycle(10) Oocyte_meiosis(8) PI3K-Akt_signaling_pathway(8) Longevity_regulating_pathway_-_worm(2) Hippo_signaling_pathway(3) Hippo_signaling_pathway_-_fly(3) Hepatitis_C(5) Hepatitis_B(3) Viral_carcinogenesis(7) |
| SGTA |  | small glutamine-rich tetratricopeptide repeat-containing protein alpha |  |
| COPB2, SEC27 |  | coatomer subunit beta' |  |
| SPN1, IWS1 |  | transcription factor SPN1 |  |
| PTPN7 |  | tyrosine-protein phosphatase non-receptor type 7 [EC:3.1.3.48] | MAPK_signaling_pathway(3) |
| TDRD9 |  | ATP-dependent RNA helicase TDRD9 [EC:3.6.4.13] |  |
| ZFP36L |  | butyrate response factor | Cellular_senescence(4) |
| ZDHHC |  | palmitoyltransferase [EC:2.3.1.225] |  |
| PXDN, VPO1 |  | peroxidase [EC:1.11.1.7] |  |
